# The Hippo network kinase STK38 contributes to protein homeostasis by inhibiting BAG3-mediated autophagy

**DOI:** 10.1101/458323

**Authors:** Christina Klimek, Ricarda Jahnke, Judith Wördehoff, Barbara Kathage, Daniela Stadel, Christian Behrends, Alexander Hergovich, Jörg Höhfeld

**Affiliations:** Institute for Cell Biology, University of Bonn, Ulrich-Haberland-Str. 61a, 53121 Bonn, Germany; Institute of Biochemistry II, Goethe University Medical School, Theodor-Stern-Kai 7, 60590 Frankfurt am Main, Germany; Munich Cluster for Systems Neurology, Ludwig-Maximilians-University Munich, Feodor-Lynen Strasse 17, 81377 München, Germany; UCL Cancer Institute, University College London, London, WC1E 6BT, UK

**Keywords:** Autophagy, proteostasis, hippo signaling, chaperone, cytoskeleton, NDR1

## Abstract

Chaperone-assisted selective autophagy (CASA) initiated by the cochaperone Bcl2-associated athanogene 3 (BAG3) represents an important mechanism for the disposal of misfolded and damaged proteins in mammalian cells. Under mechanical stress, the cochaperone cooperates with the small heat shock protein HSPB8 and the cytoskeleton-associated protein SYNPO2 to degrade force-unfolded forms of the actin-crosslinking protein filamin. This is essential for muscle maintenance in flies, fish, mice and men. Here, we identify the serine/threonine protein kinase 38 (STK38), which is part of the Hippo signaling network, as a novel interactor of BAG3. STK38 was previously shown to facilitate cytoskeleton assembly and to promote mitophagy as well as starvation and detachment induced autophagy. Significantly, our study reveals that STK38 exerts an inhibitory activity on BAG3-mediated autophagy. Inhibition relies on a disruption of the functional interplay of BAG3 with HSPB8 and SYNPO2 upon binding of STK38 to the cochaperone. Of note, STK38 attenuates CASA independently of its kinase activity, whereas previously established regulatory functions of STK38 involve target phosphorylation. The ability to exert different modes of regulation on central protein homeostasis (proteostasis) machineries apparently allows STK38 to coordinate the execution of diverse macroautophagy pathways and to balance cytoskeleton assembly and degradation.

**Abbreviations:** BAG3, BCL2-associated athanogene; CASA, chaperone-assisted selective autophagy; CHIP, carboxy terminus of HSP70 interacting protein; EPS, electrical pulse stimulation; GST, glutathione-S-transferase; mTOR, mechanistic target of rapamycin; mTORC1, mechanistic target of rapamycin complex 1; STK38, serine/threonine protein kinase 38.

## 1. Introduction

In eukaryotic cells, macroautophagy mediates the degradation of intracellular content by lysosomal hydrolytic enzymes [1,2]. It involves content enclosure by phagophore membranes, followed by fusion of the formed autophagosome with a lysosome. Autophagy was initially described as a rather unselective pathway induced under starvation conditions, when the digestion of surplus cellular material is essential for survival [3]. Since then, an increasing number of selective autophagy pathways has been identified for the regulated degradation of diverse types of cargo, including invading pathogens, dysfunctional organelles and protein aggregates [4–6]. Pathways are distinguished by the participating cargo recognition system, often involving a specialized ubiquitin conjugation machinery that labels the cargo for autophagic degradation, and by the engaged adaptor protein, which links the selected cargo to the phagophore membrane [4]. Moreover, different protein assemblies can initiate autophagosome formation such as the BECN1-RALB-exocyst complex or heteromeric complexes formed by tripartite motif (TRIM) proteins [7]. Evidently, macroautophagic degradation in eukaryotic cells does not rely on a single common machinery but involves a plethora of diverse pathways. Yet, our understanding about the separation, integration and coordination of these pathways is still very limited.

BAG3 initiates the degradation of misfolded, damaged and aggregation-prone proteins through chaperone-assisted selective autophagy (CASA) [8–11]. In neurons, BAG3-mediated autophagy contributes to the degradation of the Alzheimer-associated protein tau and pathological forms of Huntingtin and SOD1, which are causative agents of Huntington’s disease and amyotrophic lateral sclerosis, respectively [8,12–15]. In mechanically stressed cells, the pathway mediates the degradation of the actin-crosslinking protein filamin and possibly other mechanosensory proteins [10]. Under mechanical stress, filamin undergoes cycles of force-induced unfolding and refolding, which provides the structural flexibility necessary for maintaining actin contacts in this situation [16,17]. The BAG3 chaperone machinery monitors the conformational state of filamin and degrades mechanically damaged forms through CASA [18]. In agreement with a central role of BAG3 in mechanical stress protection, BAG3 was found to be essential for muscle maintenance in diverse model organisms and BAG3 impairment causes muscle dystrophy and cardiomyopathy in humans [10,19–26].

Client proteins are recognized by BAG3 in conjunction with its chaperone partners, including cytosolic members of the HSP70 chaperone family, i.e. constitutively expressed HSC70/HSPA8, and small heat shock proteins such as HSPB8 [27,28]. BAG3 promotes the ubiquitylation of chaperone-bound clients through cooperation with the HSC70 associated ubiquitin ligase STUB1/CHIP, and stimulates client sequestration in perinuclear aggregates as a prerequisite for autophagic degradation [10,13,14,29,30]. The generated ubiquitin signal is recognized by the autophagy adaptor SQSTM1/p62, which links the ubiquitylated cargo to phagophore membranes [9]. Finally, cargo enclosure can be facilitated through cooperation of BAG3 with SYNPO2 (also known as myopodin) [29]. The protein recruits a membrane fusion machinery, which mediates autophagosome formation around CASA complexes.

Of note, BAG3 exerts proteostasis functions that go beyond its involvement in autophagy [11]. In mechanically stressed cells, the cochaperone stimulates filamin transcription through engagement in the Hippo signaling pathway and increases protein synthesis by regulating the mTOR kinase [29,31]. Both measures are necessary to counteract the autophagic disposal of damaged filamin and to maintain the actin cytoskeleton under mechanical stress [11].

The Hippo pathway was first characterized as a linear kinase cascade that controls tissue growth [32]. In mammals, the pathway comprises the serine/threonine protein kinases 3 and 4 (STK3 and STK4, homologs of the *D. melanogaster* kinase Hippo) and the large tumor suppressor kinases 1 and 2 (LATS1 and LATS2). STK3/4 phosphorylate and activate LATS1/2, which in turn phosphorylate the transcriptional coactivators YAP and TAZ, causing their inactivation through cytoplasmic retention. When the pathway is switched off, for example in response to increased mechanical forces, YAP and TAZ migrate into the nucleus and stimulate the expression of target genes, including filamin [32,33]. The concept of a linear pathway, however, was recently revisited based on the identification of additional kinases that participate in Hippo signaling [34]. The serine/threonine protein kinase 38 (STK38, also known as nuclear Dbf2-related kinase 1 (NDR1)), for example, was shown to be a substrate of STK3/4 and to phosphorylate YAP [35–38]. The data place STK38 at a stage similar to that of LATS1/2 in a Hippo kinase network. Furthermore, STK38 can also be activated by STK24 [39]. This extends the network at the initiation level and provides additional means for signal input [34]. In cardiac muscle cells, STK38-mediated signaling contributes to protein homeostasis through the activation of the RNA binding protein RBM24, which mediates splicing events essential for cardiac development and for the assembly of actin-anchoring structures in this cell type [40–42].

Increasing evidence links the Hippo network to the regulation of autophagy. It was observed that STK3 and STK4 phosphorylate the autophagy membrane marker LC3, which is essential for autophagosome-lysosome fusion [43]. A prominent role in this functional context is also played by STK38. The kinase stimulates autophagosome formation in response to starvation and cell detachment by positively regulating the BECN1-RALB-exocyst complex [44]. Furthermore, STK38 participates in the initiation of mitophagy, a selective autophagy pathway for the degradation of dysfunctional mitochondria [45,46].

Strikingly, our study identifies STK38 as an inhibitor of BAG3-mediated autophagy. Interaction of the kinase with the cochaperone attenuates the autophagic degradation of the BAG3 client filamin and the BAG3 interactor SYNPO2 in adherent smooth muscle cells. Inhibition relies on a remodeling of BAG3 chaperone complexes, which involves a loss of HSPB8 and SYNPO2 binding. Despite the importance of STK38’s kinase activity for the regulation of diverse cellular processes such as proliferation or starvation induced autophagy [39,44], we found that STK38 inhibits CASA in a kinase activity independent manner. Our findings further illustrate the intensive crosstalk between the BAG3 chaperone machinery and the Hippo signaling network. Furthermore, STK38 is revealed as a cellular switch that coordinates diverse macroautophagy pathways and adjusts assembly and degradation processes during cytoskeleton homeostasis.

## 2. Material and methods

### 2.1 Antibodies

The following antibodies were used for protein detection: anti-BAG3 (rabbit; 10599-1-AP, Proteintech Group), anti-BECN1 (rabbit, #8676, Cell Signaling), anti-γ-tubulin (mouse; T5326, Sigma Aldrich), anti-filamin (rabbit; generously provided by D. Fürst, Bonn), anti-HA (rabbit, sc-805, Santa Cruz), anti-HSC70 (rabbit, generated against purified HSC70 [47]), anti-HSPB8 (rabbit, STJ24102, St Johńs Lab), anti-MBP (mouse, ab23903, Abcam), anti-SQSTM1 (guinea-pig, GP62-C, Progen), anti-RALB (rabbit, STJ28797, St Johńs Lab), anti-STK38 (mouse, ABIN564919, Abnova), anti-SYNPO2 (rabbit; generated by Dieter Fürst, Bonn) and rabbit IgG (sc-2027, Santa Cruz).

### 2.2 BAG3 interactome proteomic analysis

HEK293T human embryonic kidney cells (Sigma Aldrich, #12022001) stably expressing N-terminally HA-tagged BAG3 were harvested and lysed with 3 ml lysis buffer (50 mM Tris, pH 7.5, 150 mM NaCl, 0.5% Nonidet P40 (NP40) and EDTA-free protease inhibitor cocktail tablets). Centrifugation-cleared lysates (16,000 x g) were filtered through 0.45 µm spin filters (Millipore Ultrafree-CL) and immunoprecipitated with 60 µl anti-HA resin. Resin containing immune complexes were washed five times with lysis buffer followed by five PBS washes, and elution with 150 µl of 250 mg/ml HA peptide in PBS. Eluted immune complexes were precipitated with 20% trichloroacetic acid (TCA) and pellets were washed once with 10% TCA and four times with cold acetone. Precipitated proteins were resuspended in 50 mM ammonium bicarbonate (pH 8.0) with 20% acetonitrile and incubated with sequencing grade trypsin at a concentration of 12.5 ng/ml at 37°C for 4 h. Trypsin reactions were quenched by addition of 5% formic acid and peptides were desalted using C18 Stage Tips. For each liquid chromatography coupled to tandem mass spectrometry run using an LTQ Velos linear ion trap mass spectrometer (Thermo Scientific), 4 µl were loaded onto a 18 cm x 125 µm (ID) C18 column and peptides eluted using a 50 min 8–26% acetonitrile gradient. Spectra were acquired using a data-dependent Top-10 method. Each sample was shot twice in succession, followed by a wash with 70% acetonitrile and 30% isopropanol. Tandem MS/MS data were searched using Sequest and a concatenated target-decoy uniprot human database with a 2 Da mass window. All data were filtered to a 1% false discovery rate (peptide level) before analysis using CompPASS [48].

### 2.3 Protein expression and purification

BAG3 encoding cDNA, subcloned into pET-M11 (Novagene), was used for expression of HIS-fusion proteins in *E. coli* BL21(DE3) and purified on Ni-NTA-agarose as described by the manufacturer (Qiagen). Before elution an additional washing step with ATP was performed.

### 2.4 In-vitro binding studies

To verify binding of BAG3 to STK38, GST or GST-STK38 (#101505, Abcam) (0.5 µM) were incubated with Ni-NTA-agarose (Qiagen) in the presence or absence of equimolar amounts of Histidin-tagged BAG3 (His-BAG3). Following incubation in binding buffer (20 mM MOPS-KOH, pH 7.5, 100 mM KCl, 60 mM imidazol) for 2 h at 4°C on a rotating wheel, agarose beads were collected by centrifugation at 800 x g for 2 min and washed four times with binding buffer. Finally, bound proteins were eluted with elution buffer (20 mM MOPS-KOH, pH 7.2, 100 mM KCl, 200 mM imidazol). Eluted proteins were precipitated in 10% trichloroacetic acid overnight and analyzed by immunoblotting.

### 2.5 siRNAs and cell techniques

FlexiTube siRNAs (Qiagen) were handled according to the manufacturer. Allstars negative control siRNA (Qiagen) was used as a control. A7r5 rat smooth muscle embryonic aorta cells (Sigma Aldrich, #86050803) were cultured in Dulbecco’s Modified Eagle Medium (DMEM) containing 10% fetal calf serum, L-glutamine and penicillin/streptomycin (PS), at 37°C and 5% CO_2_. When indicated, cells were transfected with JetPRIME transfection reagent (Peqlab) following manufacturer’s instructions with plasmid DNA or siRNA. For transient transfection HeLa cells were grown to 40% confluency in a 10 cm dish. Cells were transfected with a calcium phosphate transfection method using 1.4 ml transfection solution (155 mM CaCl_2_, 137 mM NaCl, 5 mM KCl, 0.7 mM Na_2_HPO_4_, 7.5 mM glucose, 21 mM HEPES) containing 6-18 µg DNA. The transfection solution was replaced with culture medium after 16 h. Two days after transfection, cells were lysed in RIPA buffer (25 mM Tris-HCl, pH 8.0, 150 mM NaCl, 0.5% sodium deoxycholate, 1% Nonidet P-40, 0.1% SDS, 10% glycerol, complete protease inhibitor (Roche), phosSTOP phosphatase inhibitor (Roche)) and incubated on ice for 20 min. Lysates were centrifuged at 16,000 x g for 20 min at 4°C. Supernatants were collected and analyzed by immunoblotting.

To monitor SYNPO2 degradation, protein synthesis was inhibited in A7r5 cells by treatment with 50 µM cycloheximide (Roth) for indicated times prior to cell lysis. Cells were lysed as described above and protein degradation was analyzed by immunoblotting. SYNPO2 levels were quantified by densitometry and normalized to γ-tubulin levels. Level at time point zero was set to 1.

To analyze filamin degradation, A7r5 cells were seeded 48 h post transfection on fibronectin (1.25 µg/cm^2^, BD Biosciences) coated culture dishes for 24 h. Prior to cell lysis, cells were treated with 50 µM cycloheximide (Roth) for 16 h to inhibit protein synthesis (corresponding to time point zero of the experiment). Protein degradation was analyzed by immunoblotting. Filamin levels were quantified and normalized to γ-tubulin levels.

### 2.6 Immunoprecipitation

To isolate protein complexes by immunoprecipitation with an anti-FLAG (M2) antibody, HeLa cells (Sigma Aldrich, #93021013) were transfected with plasmids encoding FLAG-BAG3, FLAG-SYNPO2a, FLAG-STK38 (wild-type) and FLAG-STK38-K118R (kinase-dead variant), respectively. When indicated cells were also transfected with plasmids encoding HA-STK38, HA-STK38-K118R and HA-STK38-T444A [49]. Two days after transfection, cells were sonicated in lysis buffer (25 mM Tris-HCl, pH 8,0, 150 mM NaCl, 0.5% sodium deoxycholate, 1% Nonidet P-40, 10% glycerol, 2 mM EDTA, complete protease inhibitor (Roche)) for 20 min followed by centrifugation at 16,000 x g for 20 min at 4°C. The supernatant was adjusted to a protein concentration of 10 mg/ml. Samples were incubated with anti-FLAG M2 agarose (Sigma) (30 µl resin per 1 ml extract) for 1 h at 4°C on a rotation wheel. Agarose beads were collected by centrifugation at 800 x g for 5 min at 4°C, followed by washing four times with RIPA buffer und two times with wash buffer (20 mM MOPS pH 7.2, 100 mM KCl). When indicated, beads were incubated in ATP buffer (20 mM MOPS pH 7.2, 100 mM KCl, 2 mM MgCl_2_, 2 mM ATP) for 30 min at 37°C, prior to final elution with glycine/HCl, pH 3.5. Samples were analyzed by immunoblotting.

When complexes were immunoprecipitated with antibodies directed against SYNPO2 or BAG3, protein G-sepharose (GE Healthcare) (30 µl resin per 1 ml extract) was used together with 12.5 μg of control IgG (rabbit IgG) and the same amount of the anti-SYNPO2 or anti-BAG3 antibody.

### 2.7 Quantitative real time PCR

InviTrap Spin Universal RNA Mini Kit (Stratec) was used to isolate RNA from A7r5 rat smooth muscle cells and differentiated C2C12 myotubes. cDNA was synthesized from 1 µg RNA by using the iScript cDNA synthesis kit (Bio-Rad). Quantitative PCR (qPCR) was performed using SsoFast™ EvaGreen® Supermix (Bio-Rad). Transcript levels for GAPDH and beta-2-microglobulin (B2M) were monitored as reference transcripts. Relative gene expression was calculated using the ΔΔCT method. The specificity of the PCR amplification was verified by melting curve analysis of the final products using Bio-Rad CFX 3.1 software. For each sample three technical replicates were analyzed. The following oligonucleotide primer were used:

STK3 (*Rattus norvegicus*) forward TACGTCTATAGAAATGGCAGAAGG
STK3 (*Rattus norvegicus*) revers GATGAAAGGATGCTGTAACAGC
STK4 (*Rattus norvegicus*) forward GATCAACACGGAGGATGAGG
STK4 (*Rattus norvegicus*) revers AGCTCTTCAGAAACTCGTAGTC
STK24 (*Rattus norvegicus*) forward GACAATCGGACTCAGAAAGTGG
STK24 (*Rattus norvegicus*) revers TTCTAACAGATCCAGGGCAGAG
B2M (*Rattus norvegicus*) forward GTGATCTTTCTGGTGCTTGTC
B2M (*Rattus norvegicus*) revers AAGTTGGGCTTCCCATTCTC
GAPDH (*Rattus norvegicus*) forward GAGAAACCTGCCAAGTATGATGAC
GAPDH (*Rattus norvegicus*) revers ATCGAAGGTGGAAGAGTGGG
STK38 (*Mus musculus*) forward GAAGGAGAGGGTGACAATGAC
STK38 (*Mus musculus*) reverse GAGTCGTTTCTCTTCGTCTTTCAG
B2M (*Mus musculus*) forward CATGGCTCGCTCGGTGACC
B2M (*Mus musculus*) reverse AATGTGAGGCGGGTGGAACTG
GAPDH (*Mus musculus*) forward TGTGTCCGTCGTGGATCTGA
GAPDH (*Mus musculus*) reverse TTGCTGTTGAAGTCGCAGGAG

### 2.8 Electrical pulse stimulation

Differentiated C2C12 myotubes grown on 6-well plates were stimulated using a C-dish and the C-Pace unit (Ion Optix, Milton) for 10 msec pulses at 10 V. A pulse program of 5 sec at 15 Hz (tetanic hold), 5 sec pause, 5 sec at 5 Hz (twitch contractions) and 5 sec pause was repeated, for indicated durations, as described previously [50]. Myotubes were harvested in PBS and lysed as described above. Protein level was analyzed by immunoblotting.

### 2.9 Statistics

Quantification of blot lane intensities was accomplished using ImageJ. Data were analyzed for statistical significance using the two-tailed student’s t-test. P-value levels derived from student’s t-test are shown with significance stars: *p ≤ 0.05, **p ≤ 0.01, ***p ≤ 0.001.

## 3. Results

### 3.1 STK38 interacts with BAG3

BAG3 initiates the degradation of misfolded, damaged and aggregation-prone proteins through CASA by interacting with diverse partner proteins (Fig. 1A) [10,11,13,14,27–29,31]. To further elucidate the BAG3 interactome, an hemagglutinin (HA) tagged form of the cochaperone was expressed in human HEK293T cells followed by BAG3 complex isolation and mass spectrometry, according to previously established procedures [51]. Analysis of complex composition revealed a substantial overlap with the BAG3 interactome deduced from the BioGRID protein interaction database (Fig. S1, Supplemental Table S1, BAG3 BioGRID entry). Because of the functional interplay of BAG3 with the Hippo signaling network, we screened the data set, including low scoring potential interactors, for the presence of Hippo network components [52] (Supplemental Table S1, Hippo network). Among BAG3-associated proteins was the kinase STK38, a recent addition to the Hippo network [34,53] (Supplemental Table S1, highlighted in yellow). STK38 has been shown to regulate cytoskeleton assembly and diverse macroautophagy pathways [42,44–46,54]. We therefore decided to further investigate the interaction and possible functional interplay of STK38 with BAG3.

**Fig. 1.**
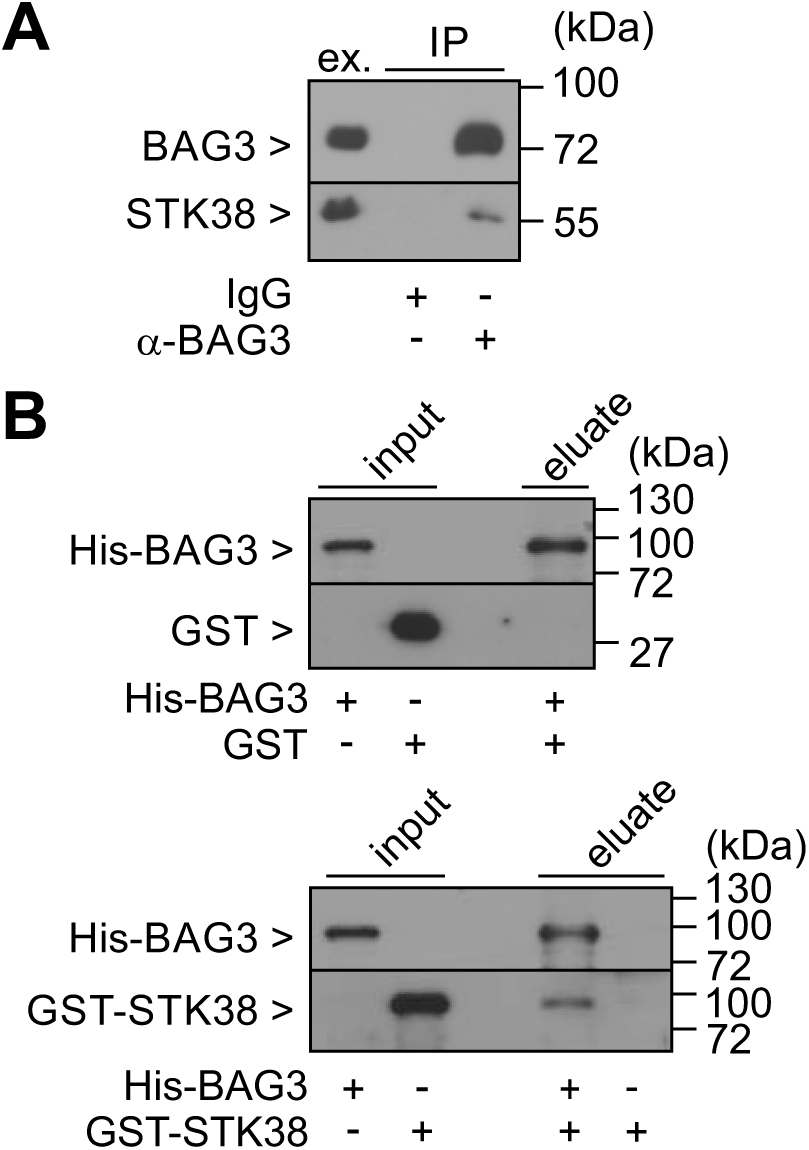
The CASA-mediating cochaperone BAG3 interacts with STK38. (A) STK38 was detectable in endogenous BAG3 complexes isolated from A7r5 cells by immunoprecipitation (IP) with an anti-BAG3 antibody (α-BAG3). A control sample received unrelated rabbit IgGs (IgG). Ex. corresponds to 40 µg protein of the cell extract. Indicated proteins were detected by western blotting with corresponding antibodies. (B) BAG3 directly interacts with STK38. Purified histidine-tagged BAG3 (His-BAG3) was incubated at equimolar concentration with 0.5 µM GST or a GST-STK38 fusion protein, followed by affinity chromatography on Ni-NTA agarose. Input corresponds to 2% of the incubated proteins, the eluate represents 30% of the total sample. Indicated proteins were detected with an anti-GST and anti-BAG3 antibody, respectively. (See also Supplemental Figure S1 and Table S1.)

As a first step, the observed interaction was verified for endogenous proteins. An anti-BAG3 antibody was used for immunoprecipitation of endogenous BAG3 complexes from A7r5 rat smooth muscle cells, which were subsequently probed for the presence of STK38 (Fig. 1A). The kinase was readily detectable in the isolated BAG3 complexes, pointing to a physiologically relevant interaction.

To investigate whether STK38 interacts directly with BAG3, in-vitro binding studies were performed with purified proteins. Histidine-tagged BAG3 (His-BAG3) was incubated at equimolar concentration (0.5 µM) with glutathione-S-transferase (GST) or a GST-STK38 fusion protein followed by affinity chromatography on Ni-NTA agarose. While GST was not detectable in association with BAG3, GST-STK38 was retained on the affinity resin in the presence of BAG3 and did not bind to the resin when the cochaperone was omitted (Fig. 1B). The data reveal a direct interaction between STK38 and BAG3.

### 3.2 STK38 inhibits CASA

To analyze functional consequences of the STK38-BAG3 interaction, the CASA-mediated turnover of the actin-crosslinking protein filamin was investigated. A7r5 rat smooth muscle cells were grown on the extracellular matrix protein fibronectin to induce mechanical tension within the actin cytoskeleton based on cell-matrix contacts. Under these conditions, ∼ 40% of the cellular filamin was degraded during inhibition of protein synthesis by cycloheximide for 16 hours (Fig. 2A). This is consistent with previous findings, which showed that degradation is dependent on BAG3 and SYNPO2 and requires inhibition of the mTOR kinase [29,31]. To verify the impact of STK38 on filamin turnover, the kinase was depleted in A7r5 cells by siRNA transfection (Fig. 2B). Of note, STK38 depletion significantly stimulated filamin degradation, leading to a reduction by about 60% (Fig. 2A). This finding points to an inhibitory role of STK38 in the regulation of BAG3-mediated autophagy. In further support of this conclusion, STK38 depletion also reduced the levels of the autophagic ubiquitin adaptor SQSTM1 upon cycloheximide treatment (Fig. 2A). SQSTM1 facilitates cargo loading and enclosure during BAG3-mediated autophagy [9].

**Fig. 2.**
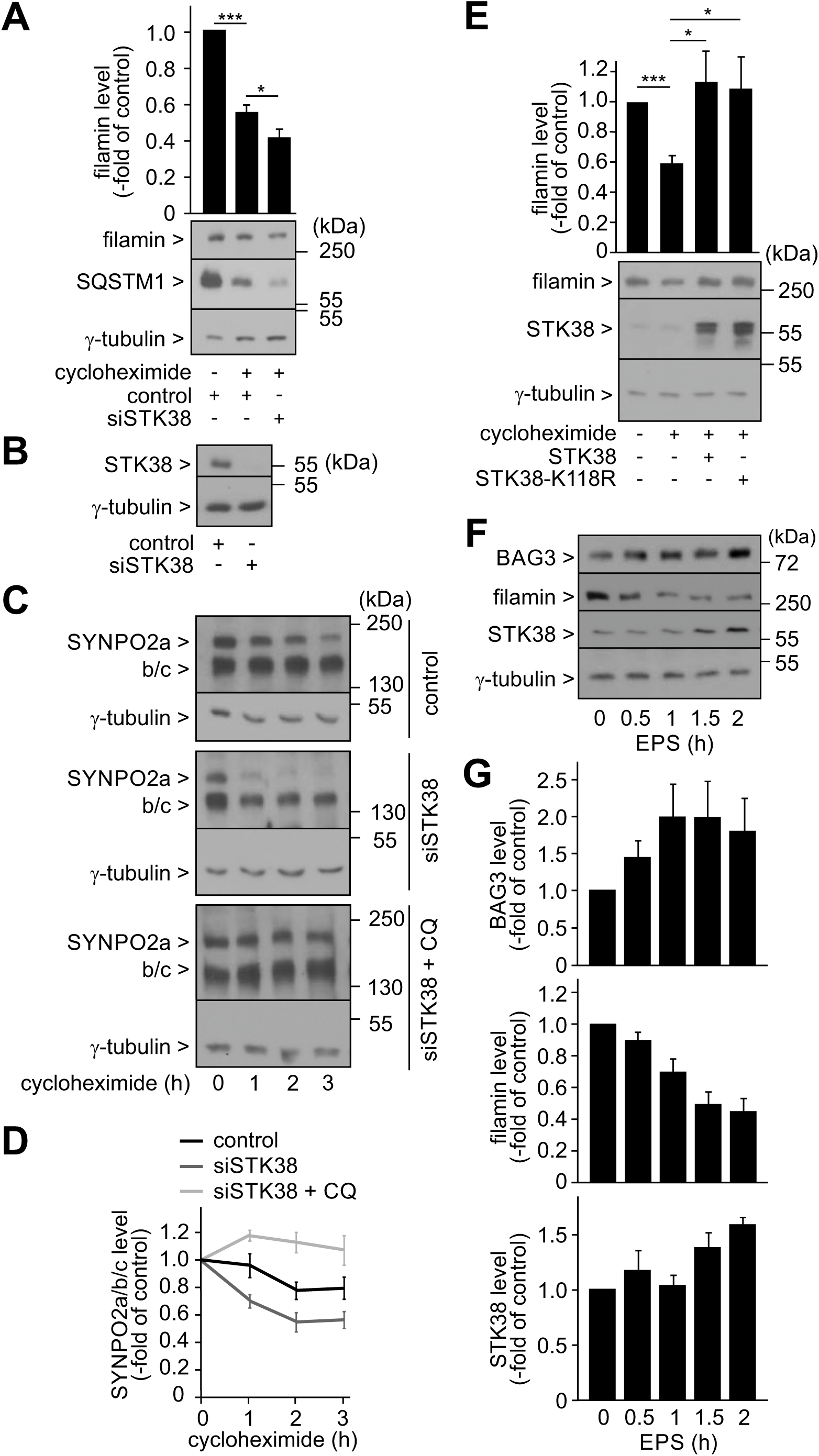
STK38 inhibits BAG3-mediated autophagy. (A) Depletion of STK38 facilitates the degradation of filamin. A7r5 cells were transiently transfected for 72 h with siRNA directed against STK38 (siSTK38). Control cells received allstars negative control siRNA (control). 24 h before lysis cells were transferred to fibronectin-coated culture dishes. When indicated, cells were incubated with cycloheximide (50 µM) for 16 h. 50 µg protein were loaded per lane. Filamin, SQSTM1 and γ-tubulin were detected by western blotting with corresponding antibodies. Signal intensities were quantified by densitometry, whereby the filamin level was normalized to the level of γ-tubulin detected in the same sample. Filamin level in control cells was set to 1. Data represent mean values +/- SEM: n ≥ 16, *p ≤ 0.05, ***p ≤ 0.001. (B) STK38 was depleted in A7r5 cells transiently transfected with a corresponding siRNA (siSTK38) for 72 h. Control cells received allstars negative control siRNA (control). 50 µg protein were loaded per lane. Indicated proteins were detected by western blotting with corresponding antibodies. (C) Depletion of STK38 stimulates the autophagic degradation of SYNPO2 in adherent A7r5 cells. A7r5 cells were transiently transfected for 48 h with siRNA directed against STK38 (siSTK38). Control cells received allstars negative control siRNA (control). Treatment with chloroquine (100 µM) was performed for 16 h prior to lysis (CQ). When indicated, cells were incubated with cycloheximide (50 µM) for 1, 2 or 3 h before lysis. 50 µg protein were loaded per lane. Isoforms a, b and c of SYNPO2 and γ-tubulin were detected by western blotting with corresponding antibodies. Signal intensities were quantified, whereby the level of SYNPO2 isoforms was normalized to the level of γ-tubulin detected in the same sample. (D) Quantification of the data presented in (C). Level of SYNPO2 isoforms detected in control cells at time point 0 was set to 1. Data represent mean values +/- SEM: n ≥ 5. (E) Overexpression of STK38 and the kinase-dead variant STK38-K118R inhibits the degradation of filamin in adherent smooth muscle cells. A7r5 cells were transiently transfected with plasmids encoding STK38 and STK38-K118R for 72 h. Control cells received empty vector. 24 h before lysis cells were transferred to fibronectin-coated culture dishes. When indicated, cells were incubated with cycloheximide (50 µM) for 16 h. 50 µg protein were loaded per lane. Filamin, STK38 and γ-tubulin were detected by western blotting with corresponding antibodies. Signal intensities were quantified, whereby the filamin level was normalized to the level of γ-tubulin detected in the same sample or to the protein amount determined by SDS-PAGE and Ponceau S staining. Filamin level in control cells was set to 1. Data represent mean values +/- SEM: n ≥ 15, *p ≤ 0.05, **p ≤ 0.01, ***p ≤ 0.001. (F) STK38 expression is induced in murine myotubes subjected to prolonged mechanical stress and induction correlates with an attenuation of filamin degradation. Differentiated C2C12 myotubes were subjected to electrical pulse stimulation (EPS) for the indicated times, followed by lysis and western blotting. 60 µg protein were loaded per lane. Indicated proteins were detected using specific antibodies. (G) Quantification of the data presented in (F). Level of indicated proteins detected at time point 0 was set to 1. Data represent mean values +/- SEM: n ≥ 4.

We also investigated the degradation of SYNPO2. Isoforms of SYNPO2, which possess an amino-terminal PDZ domain, i.e. SYNPO2a, b and c (isoforms b and c cannot be distinguished by SDS-PAGE because of similar molecular weight, Fig. 2C), cooperate with BAG3 to initiate autophagy and are co-degraded during the process [29]. SYNPO2 turnover can therefore be used as a readout for CASA activity. Depletion of STK38 did not affect the steady state levels of SYNPO2 (Fig. S2, compare lanes 1 and 3). Impact on SYNPO2 degradation might be masked in this situation by altered synthesis. To monitor SYNPO2 turnover, we therefore also performed cycloheximide shut-off experiments. (Fig. 2C and D). Treatment of adherent A7r5 smooth muscle with cycloheximide for three hours led to a significant reduction of SYNPO2 levels (Fig. S2 and Fig. 2C). Notably, depletion of STK38 resulted in a further decrease of SYNPO2 levels, consistent with increased degradation in the absence of the Hippo network kinase (Fig. 2C and D). Moreover, treatment with chloroquine (CQ), which inhibits CASA and other autophagic degradation pathways, attenuated SYNPO2 degradation in STK38 depleted cells (Fig. 2C and D). The data further support an inhibitory activity of STK38 in CASA regulation.

To exclude off-target effects of the STK38 siRNA, a complementation assay was performed. Adherent A7r5 cells were transfected with the siRNA directed against the rat transcript together with or without a plasmid encoding human STK38. Cycloheximide treatment revealed again increased turnover of SYNPO2 isoforms upon STK38 depletion (Fig. S2B). Importantly, SYNPO2 degradation was blocked when siRNA transfected cells also received the overexpression plasmid (Fig. S2B). This excludes any off-target effects of the STK38 siRNA and illustrates that CASA activity is inversely correlated with the expression level of STK38 in adherent smooth muscle cells.

Furthermore, we observed that overexpression of STK38 blocked the degradation of filamin, in agreement with a role of STK38 as a CASA inhibitor (Fig. 2E). Besides the plasmid for overexpression of wild-type STK38, we also used one encoding a variant of STK38 that completely lacks any kinase activity [49,55,56]. In this variant (STK38-K118R) a lysine at position 118, which is essential for ATP binding, is replaced by arginine, thereby abolishing kinase activity. Notably, this kinase-dead variant rescued SYNPO2 turnover in the complementation assay (Fig. S2B) and stabilized filamin like wild-type STK38 (Fig. 2E). The findings indicate that STK38 inhibits CASA in a kinase activity independent manner.

### 3.3 STK38 expression and CASA activity are regulated under mechanical stress

CASA is induced in mechanically stressed cells and tissues, when cytoskeleton components become unfolded and are damaged by mechanical force [29,57]. Cytoskeleton contraction and damage were triggered in differentiated murine C2C12 myotubes through electrical pulse stimulation (EPS) according to previously established protocols [50]. Subsequently, mechanical stress dependent changes in protein abundance were monitored. EPS caused an immediate increase in BAG3 expression and a rapid decline of filamin levels in mechanically stressed myotubes during the first hour of treatment, reflecting force-induced CASA activity (Fig. 2F and G). At subsequent time points, however, filamin levels remained constant despite elevated BAG3 expression. Strikingly, the attenuation of filamin degradation coincided with an increase of STK38 levels at late time points of EPS (Fig. 2F and G). STK38 upregulation may thus limit CASA activity upon severe and prolonged mechanical stress, when over-excessive degradation of cytoskeleton components would contribute to a collapse of the cellular architecture, and may initiate repair processes based on an STK38-dependent stimulation of cytoskeleton assembly [54].

Quantitative PCR revealed a significant upregulation of STK38 transcription between 0.5 and 1 h of EPS stimulation, which precedes the accumulation at the protein level (Fig. S3). The data illustrate that STK38 expression is regulated by mechanical stimuli in smooth muscle cells.

### 3.4 The STK38 upstream kinases STK3, STK4 and STK24 participate in CASA regulation

STK38 is part of the Hippo signaling network [32]. STK3 and STK4 act upstream of STK38 and are able to activate the kinase [35,36,38]. Furthermore, the kinase STK24 can positively regulate STK38 through phosphorylation [39]. We investigated whether STK3, STK4 and STK24 participate in the regulation of CASA. Expression of the kinases was reduced by transient transfection of A7r5 cells with corresponding siRNAs, as verified through transcript quantification (Fig. 3A). To monitor consequences for SYNPO2 degradation, cycloheximide shut-off experiments were performed (Fig. 3B and C). Depletion of STK3 or STK4 attenuated the degradation of SYNPO2 during cycloheximide treatment, indicating that these kinases are required for the execution of CASA (Fig. 3B). Notably, STK3 and STK4 were previously identified as essential autophagy regulators, which mediate the phosphorylation of the autophagosome membrane marker LC3 to facilitate autophagosome-lysosome fusion [43]. The observed inhibition of SYNPO2 degradation may thus reflect a general requirement of STK3 and STK4 for macroautophagic degradation.

**Fig. 3.**
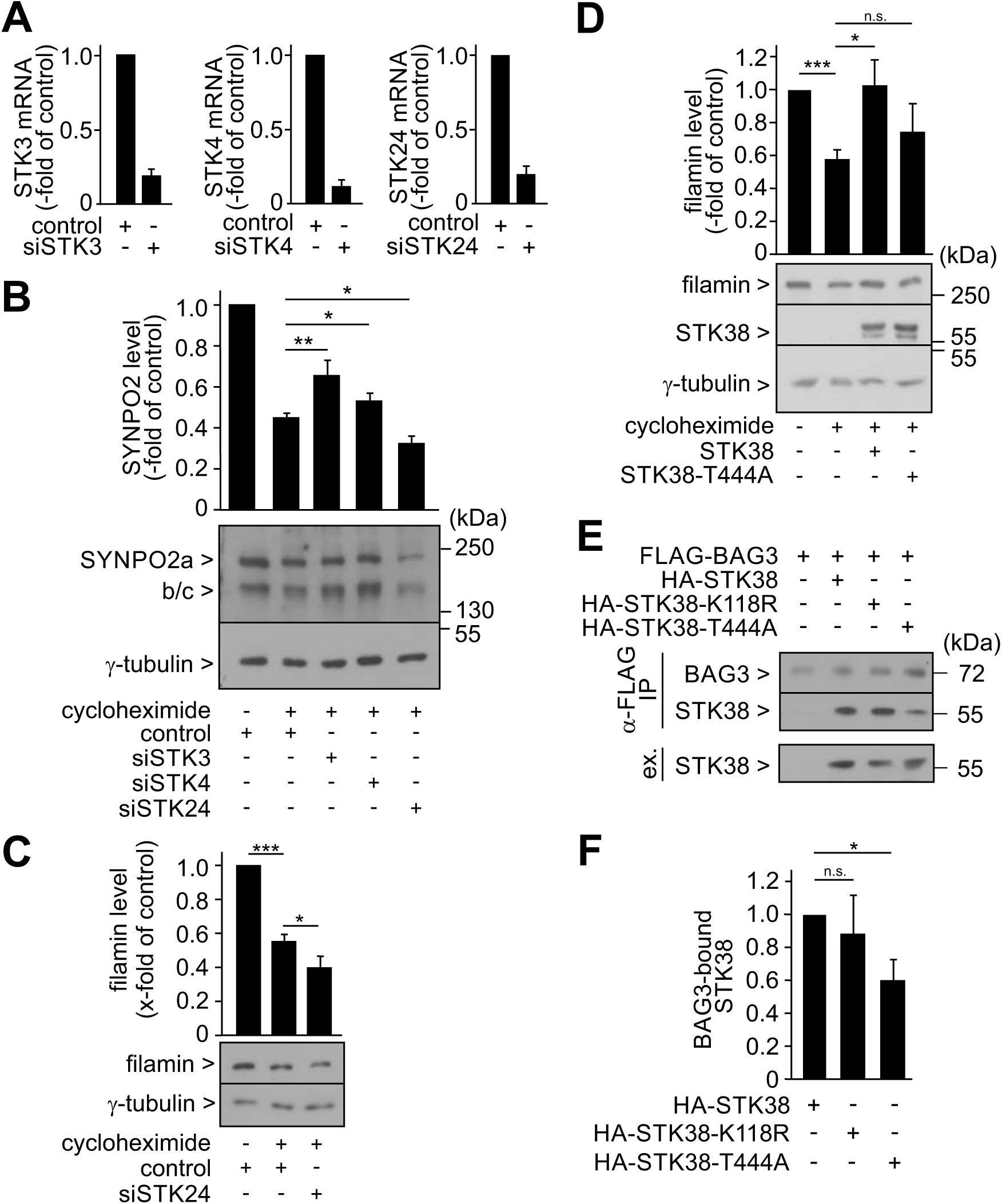
Hippo kinases that act upstream of STK38 contribute to the regulation of CASA. (A) A7r5 cells were transiently transfected with siRNAs directed against STK3, STK4 and STK24 for 48 h. Control cells received allstars negative control siRNA (control). Transcript levels were determined by quantitative real time PCR. Data represent mean values +/- SEM: n = 3. (B) Depletion of STK3 and STK4 attenuates SYNPO2 degradation, whereas depletion of STK24 facilitates turnover. A7r5 cells were transiently transfected for 48 h with siRNA directed against the indicated kinases. Control cells received allstars negative control siRNA (control). When indicated, cells were incubated with cycloheximide (50 µM) for 3 h before lysis. 60 µg protein were loaded per lane. Isoforms a, b and c of SYNPO2 and γ-tubulin were detected by western blotting with corresponding antibodies. Signal intensities were quantified and normalized to the level of γ-tubulin or total protein detected in the same sample. SYNPO2 level in control cells, which did not receive cycloheximide, was set to 1. Data represent mean values +/- SEM: n ≥ 10, *p ≤ 0.05, **p ≤ 0.01. (C) Depletion of STK24 facilitates the degradation of filamin. A7r5 cells were transiently transfected for 72 h with siRNA directed against STK24 (siSTK24) or control siRNA (control). 24 h before lysis cells were transferred to fibronectin-coated culture dishes. When indicated, cells were incubated with cycloheximide (50 µM) for 16 h. 50 µg protein were loaded per lane. Filamin and γ-tubulin were detected by western blotting with corresponding antibodies. Signal intensities were quantified by densitometry, whereby the filamin level was normalized to the level of γ-tubulin detected in the same sample. Filamin level in control cells was set to 1. Data represent mean values +/- SEM: n ≥ 15, *p ≤ 0.05, ***p ≤ 0.001. (D) Overexpression of STK38-T444A, which cannot be phosphorylated by STK38 upstream kinases, does not significantly affect filamin degradation. A7r5 cells were transiently transfected with plasmids encoding STK38 and STK38-T444A for 72 h. Control cells received empty vector. 24 h before lysis cells were transferred to fibronectin-coated culture dishes. When indicated, cells were incubated with cycloheximide (50 µM) for 16 h. 50 µg protein were loaded per lane. Filamin, STK38 and γ-tubulin were detected by western blotting with corresponding antibodies. Signal intensities were quantified, whereby the filamin level was normalized to the level of γ-tubulin detected in the same sample. Filamin level in control cells was set to 1. Data represent mean values +/- SEM: n ≥ 6, *p ≤ 0.05, **p ≤ 0.01, ***p ≤ 0.001. (E) STK38-T444A displays reduced affinity for BAG3. HeLa cells were transiently transfected with plasmids encoding FLAG-BAG3, HA-STK38, HA-STK38-K118R and HA-STK38-T444A as indicated for 48 h, followed by immunoprecipitation with an anti-FLAG antibody resin (α-FLAG IP). Immunoprecipitated complexes were analyzed by western blotting using specific antibodies. Ex. corresponds to 50 µg protein of the cell lysates. (F) Quantification of data obtained under (E). STK38 level detected in BAG3 complexes was set to 1. Data represent mean values +/- SEM: n ≥ 5, *p ≤ 0.05, **p ≤ 0.01, ***p ≤ 0.001.

In contrast to findings for STK3 and STK4, depletion of STK24 stimulated the turnover of SYNPO2 in adherent A7r5 cells (Fig. 3B). Moreover, also filamin degradation was increased upon STK24 depletion (Fig. 3C). Thus, the data for STK24 mirror those obtained for STK38 (see above). Our findings suggest that STK24 cooperates with STK38 on a signaling pathway that limits CASA activity in mammalian cells.

STK24 phosphorylates STK38 on threonine 444 (T444) [39]. A mutant form of STK38, which cannot be phosphorylated due to a threonine to alanine exchange at this position (T444A), was used to explore the importance of STK38 regulation for CASA inhibition. In contrast to wild-type STK38, the mutant form did not significantly attenuate filamin degradation upon overexpression in adherent A7r5 smooth muscle cells (Fig. 3D). Because T444 phosphorylation could affect binding of STK38 to BAG3, association between both proteins was analyzed. HeLa cells were transiently transfected with plasmid constructs encoding epitope-tagged forms of BAG3 and STK38, STK38-K118R or STK38-T444A, followed by BAG3 complex isolation and detection of STK38 variants in these complexes (Fig. 3E). Similar levels of wild-type STK38 and the kinase-dead variant K118R were found associated with immunoprecipitated BAG3 (Fig. 3E and F). However, the T444A variant, although expressed at the same level as STK38 and STK38-K118R, was recovered to a lesser extent in BAG3 complexes compared to the other variants (Fig. 3E and F). Taken together, our data suggest that phosphorylation of STK38 at threonine 444 by upstream kinases such as STK24 promotes the association of STK38 with BAG3 for efficient inhibition of CASA activity.

### 3.5 CASA is initiated independently of the STK38 targets BECN1 and RALB

During starvation and detachment induced autophagy, STK38 interacts with the autophagy regulator BECN1 to promote its association with RALB and the exocyst membrane tethering complex at initial stages of autophagosome formation [44]. This stimulating function, which is dependent on the kinase activity of STK38 [44,46], is in sharp contrast to the inhibitory and kinase activity independent function exerted by STK38 in the context of CASA (see Fig. 2). Autophagosome membrane formation and expansion during CASA may thus mechanistically differ from early stages of the other autophagy pathways. To verify this, BECN1 and RALB were depleted from A7r5 smooth muscle cells (Fig. 4A), and consequences for CASA activity were investigated by monitoring the turnover of SYNPO2 and filamin in cycloheximide shut-off experiments. Notably, BECN1 and RALB depletion, respectively, did not have any significant effect on the degradation of SYNPO2 or filamin (Fig. 4B and C). CASA apparently proceeds in a BECN1 and RALB independent manner, providing a molecular basis for the different mode of regulation exerted by STK38 on CASA in comparison to other macroautophagy pathways.

**Fig. 4.**
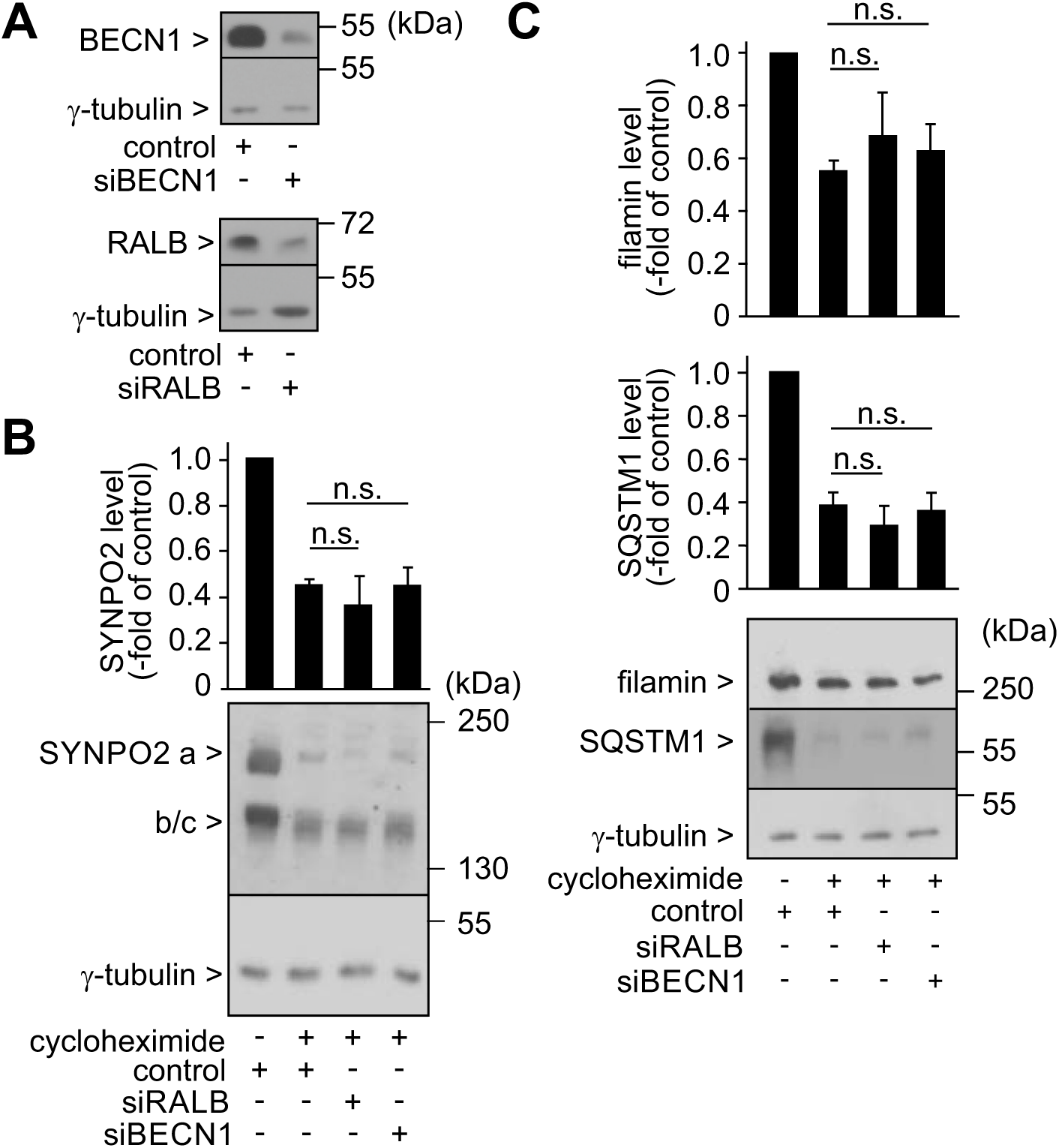
CASA proceeds independently of BECN1 and RALB. (A) BECN1 and RALB were depleted in A7r5 cells transiently transfected with corresponding siRNAs for 72 h. Control cells received allstars negative control siRNA (control). 50 µg protein were loaded per lane. Indicated proteins were detected by western blotting with corresponding antibodies. (B) SYNPO2 degradation is not affected by depletion of BECN1 or RALB in adherent smooth muscle cells. A7r5 cells were transiently transfected for 72 h with siRNA directed against the indicated proteins. Control cells received allstars negative control siRNA (control). When indicated, cells were incubated with cycloheximide (50 µM) for 4 h before lysis. 60 µg protein were loaded per lane. SYNPO2 isoforms and γ-tubulin were detected by western blotting with corresponding antibodies. Signal intensities were quantified, whereby the level of SYNPO2 isoforms was normalized to the level of γ-tubulin detected in the same sample. SYNPO2 level in control cells, which did not receive cycloheximide, was set to 1. Data represent mean values +/- SEM: n ≥ 4. (n.s. - not significant). (C) Depletion of BECN1 and RALB does not affect the degradation of filamin and SQSTM1 in adherent A7r5 cells. A7r5 cells were transiently transfected for 72 h with siRNA directed against RALB (siRALB), BECN1 (siBECN1) or control siRNA (control). 24 h before lysis cells were transferred to fibronectin-coated culture dishes. When indicated, cells were incubated with cycloheximide (50 µM) for 16 h. 50 µg protein were loaded per lane. Filamin, SQSTM1 and γ-tubulin were detected by western blotting with corresponding antibodies. Signal intensities were quantified by densitometry, whereby the filamin level was normalized to the level of γ-tubulin detected in the same sample. Filamin level in control cells was set to 1. Data represent mean values +/- SEM: n ≥ 13 (n.s. - not significant).

We also noted that degradation of the autophagic ubiquitin adaptor SQSTM1 was not altered upon BECN1 and RALB depletion (Fig. 4C), suggesting that CASA is the prevalent macroautophagy pathway in adherent smooth muscle cells.

### 3.6 STK38 disrupts the interaction of BAG3 with HSPB8 and SYNPO2

Next, we sought to elucidate the molecular basis for the inhibitory function of STK38 in CASA regulation. In particular, the impact of STK38 on the composition of BAG3 chaperone complexes was investigated. For this purpose, STK38 and BAG3 were transiently overexpressed in HeLa cells, followed by immunoprecipitation of BAG3 complexes (Fig. 5A). Purified matrix-bound complexes were incubated with ATP, which disrupts the interaction of BAG3 with the ATP-regulated chaperone HSC70 and causes partial release of non-antibody bound BAG3 from oligomeric chaperone complexes (Fig. 5A). Binding of BAG3 to the small heat shock protein HSPB8 is not ATP-sensitive [10,27]. Indeed, HSPB8 was only detectable in eluates obtained after glycine-induced disruption of the antibody-antigen interaction (Fig. 5A). Most notably, the presence of STK38 in BAG3 complexes had a profound impact on the protein composition of these complexes. While association of the cochaperone with HSC70 was not affected, when STK38 entered BAG3 complexes, it strongly reduced the interaction of HSPB8 with BAG3 (Fig. 5A).

**Fig. 5.**
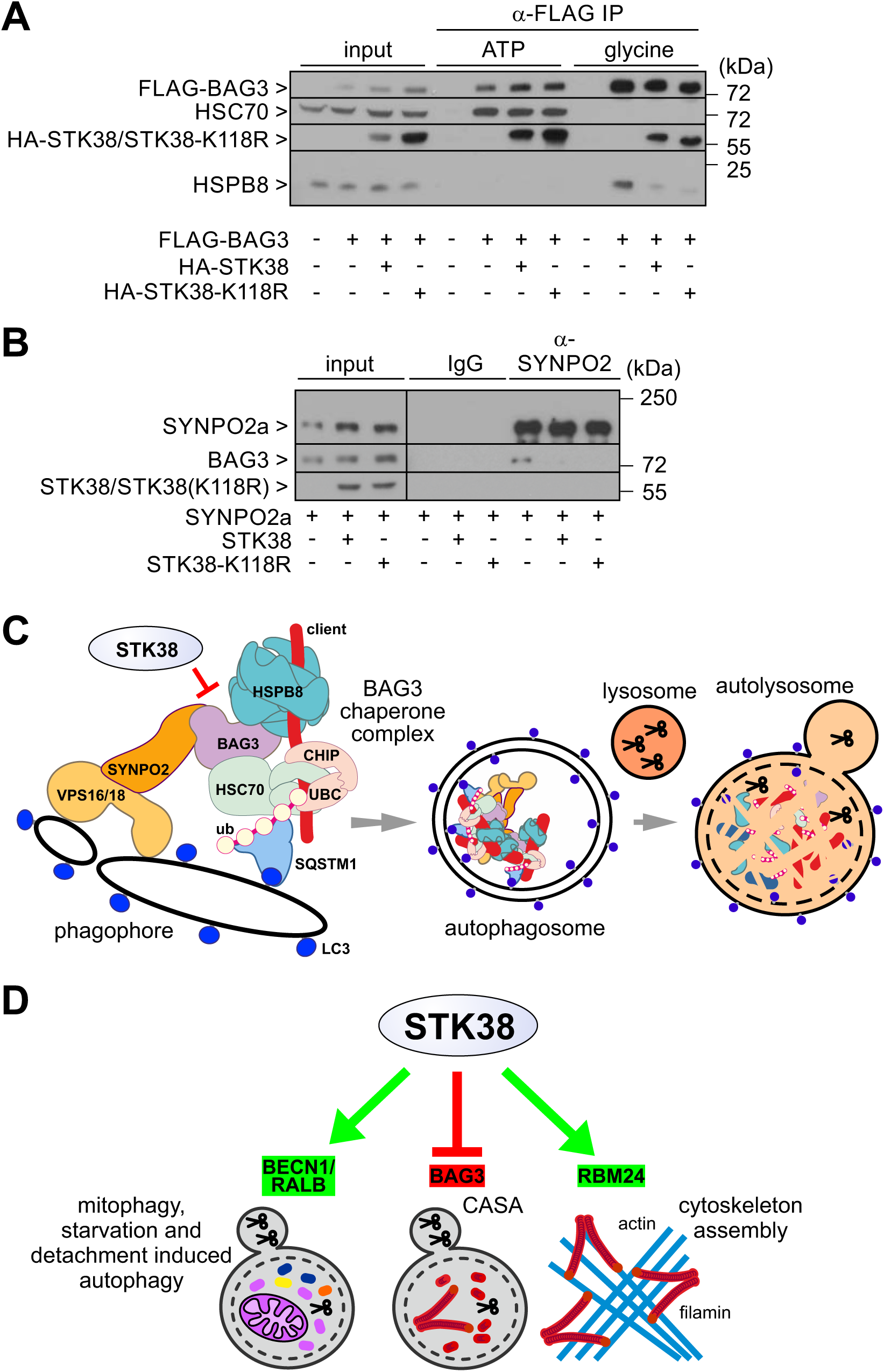
STK38 disrupts the interaction of BAG3 with HSPB8 and SYNPO2 in a manner independent of its kinase activity. (A) Association of STK38 with BAG3 interferes with the binding of the cochaperone to HSPB8. HeLa cells were transiently transfected with plasmids encoding FLAG-BAG3, HA-STK38 and HA-STK38-K118R as indicated for 48 h, followed by immunoprecipitation with an anti-FLAG antibody resin (α-FLAG IP). Isolated complexes were treated with ATP (ATP) prior to elution with glycine containing buffer (glycine). Indicated proteins were detected by western blotting using specific antibodies. Input corresponds to 50 µg protein of the cell lysates. (B) Overexpression of STK38 or the kinase-dead variant STK38-K118R in A7r5 cells disrupts the interaction of SYNPO2a with BAG3. HeLa cells were transiently transfected with plasmids encoding SYNPO2a, STK38 and STK38-K118R as indicated for 48 h, followed by immunoprecipitation with an anti-SYNPO2 antibody (α-SYNPO2). Control samples received unrelated rabbit IgGs (IgG) instead. Indicated proteins were detected by western blotting using specific antibodies. Input corresponds to 50 µg protein of the cell lysates. (C) Schematic presentation of the CASA pathway. (UBC - ubiquitin conjugating enzyme, ub - ubiquitin) (D) STK38 acts as a central switch during protein homeostasis in mammalian cells. It positively regulates mitophagy and starvation and detachment induced autophagy through BECN1 and RALB, whereas STK38 inhibits CASA through association with BAG3. Stimulation of cytoskeleton assembly relies on the activation of RBM24 by STK38.

STK38 exerts its CASA inhibiting activity in a kinase independent manner (see Fig. 2E). Therefore, we also investigated whether the kinase-dead variant of STK38 (STK38-K118R) affects BAG3 complex composition. As is evident from figure 5A, association of STK38-K118R with BAG3 disrupted the interaction of the cochaperone with HSPB8. The kinase-dead variant thus behaves like the wild-type protein, further demonstrating the kinase activity independent mode of regulation that is exerted by STK38 on BAG3.

We had previously observed that SYNPO2 could not be detected in BAG3 complexes isolated by immunoprecipitation [29]. To verify, whether STK38 also affects the BAG3-SYNPO2 interaction, SYNPO2 was transiently overexpressed in HeLa cells alone or together with STK38 or the kinase-dead variant, and an anti-SYNPO2 antibody was subsequently used for complex isolation by immunoprecipitation. While BAG3 was associated with SYNPO2 under control conditions, the cochaperone was not detectable in SYNPO2 complexes upon overexpression of STK38 or STK38-K118R (Fig. 5B). Moreover, neither variant of STK38 could be detected in association with SYNPO2. Binding of STK38 or STK38-K118R to BAG3 apparently interferes with the interaction of the cochaperone with SYNPO2. Taken together, our experiments reveal a kinase-independent ability of STK38 to remodel BAG3 chaperone complexes, leading to a loss of HSPB8 and SYNPO2 (Fig. 5C). Because both proteins cooperate with BAG3 during CASA (Fig. 5C) [8,10,29,30], STK38 binding to BAG3 potently interferes with the execution of the macroautophagic degradation pathway.

## 4. Discussion

The Hippo network kinase STK38 is essential for cytoskeleton assembly and for the execution of mitophagy as well as starvation and detachment induced autophagy [44–46]. Here, we demonstrate that binding of STK38 to the cochaperone BAG3 inhibits the degradation of cytoskeleton proteins through chaperone-assisted selective autophagy. Taken together, it becomes apparent that STK38 coordinates the execution of diverse macroautophagy pathways and controls cytoskeleton maintenance in mammalian cells (Fig. 5D).

During starvation and detachment induced autophagy, STK38 promotes the formation of the BECN1-RALB-exocyst complex [44]. The complex activates the phosphatidylinositol-3 kinase VPS34, which generates PI3P-positive precursor membranes for the recruitment of additional autophagy factors at initial stages of autophagosome formation [2,58]. STK38 also stimulates mitophagy, the autophagic clearance of damaged and depolarized mitochondria [45,46]. Mitophagy is triggered by the exposure of ubiquitin chains on the mitochondrial outer membrane, generated by the ubiquitin ligase Parkin [59]. Targeting of Parkin to depolarized mitochondria is facilitated by STK38 [45]. Furthermore, the ability of STK38 to stimulate the formation of the BECN1-RALB-exocyst complex may contribute to the induction of mitophagy, as RALB depletion attenuates STK38-dependent mitophagy in transformed human cells [46]. Importantly, we demonstrate here that CASA proceeds independently of BECN1 and RALB (Fig. 4). This is consistent with a previous study showing that BECN1 is not required for CASA [60]. The fact that components, which confer the stimulating activity of STK38 during mitophagy and starvation and detachment induced autophagy, are dispensable for CASA, apparently provides the basis for a different mode of regulation, enabling STK38 to inhibit CASA (Fig. 5C and D).

Surprisingly, CASA inhibition did not require the kinase activity of STK38. The kinase-dead variant STK38-K118R [49,55,56] attenuated filamin and SYNPO2 turnover like the wild-type protein upon overexpression in adherent smooth muscle cells (Figs. 2C and S2B). Moreover, binding of both forms to BAG3 interfered with the interaction of the cochaperone with HSPB8 and SYNPO2 (Fig. 5A and B). Thus, steric hindrances rather than phosphorylation-induced conformational changes seem to cause the STK38-induced disruption of BAG3 complexes and CASA inhibition.

Although independent of the kinase activity of STK38, CASA inhibition nevertheless appears to be a regulated event. The STK38 upstream kinase STK24 [39] also attenuated CASA (Fig. 4), and a mutant form of STK38 that cannot be activated by STK24 (STK38-T444A) displayed reduced BAG3 binding and CASA inhibiting activity (Fig. 4). STK24 and STK38 thus seem to cooperate on a signaling pathway, which limits autophagic flux through CASA in mammalian cells.

Mechanical force is a major physiological signal in the regulation of the Hippo network and CASA activity [11,61]. Intriguingly, regulation is inversely correlated. Force generated in adherent cells or contracting muscles stimulates CASA [29,57], whereas the loss of mechanical cues for example upon cell detachment from the extracellular matrix activates Hippo kinases [33,62]. In this regard, the observed STK38-mediated regulation of BAG3 seems to represent an important nodal point for co-regulation. Through STK38 and the STK38 upstream kinase STK24, a detachment-induced activation of Hippo kinases would be transferred onto the CASA machinery to switch off the degradation pathway, no longer needed for the disposal of force-unfolded cytoskeleton proteins in this situation. At the same time, core macroautophagy factors would become available for detachment-induced autophagy and mitophagy, which are positively regulated by STK38 (Fig. 5D) [46,63]. The autophagic degradation of mechanotransducers and adhesion complexes prevents cell death, when cells are deprived of extra cellular matrix contacts, and mitophagy would limit the production of reactive oxygen species in this situation [46,64–66]. In a similar manner, STK38-mediated regulation could operate upon nutrient depletion and redirect common macroautophagy factors towards starvation-induced autophagy to ensure survival at the expense of CASA and cytoskeleton quality control.

Importantly, functional cooperation between the Hippo network and the BAG3 chaperone machinery is not a one-way route, on which Hippo signaling controls CASA activity. We previously showed that BAG3 directly interacts with the Hippo kinases LATS1 and LATS2 in a manner that disrupts inhibition of the transcriptional coactivators YAP and TAZ [29]. The study revealed a BAG3-dependent mechanotransduction pathway that controls downstream components of the Hippo network. On this pathway, accumulation of force-unfolded proteins triggers the activation of the heat shock transcription factor 1 (HSF1), which stimulates BAG3 expression and ultimately YAP/TAZ activation [29]. Thus, bi-directional crosstalk connects the Hippo network and the BAG3 chaperone machinery.

In striated skeletal and heart muscle, BAG3 is localized at actin-anchoring Z-disks, which limit the contractile sarcomeric unit [67]. Loss or functional impairment of the cochaperone in animal models and patients results in a force-induced disintegration of Z-disks and aggregation of Z-disk proteins, leading to severe muscle weakness [10,19,20,22,24,68,69]. Intriguingly, knockdown of STK38 in cardiac muscle cells essentially mirrors the phenotype caused by BAG3 depletion or impairment. It also triggers Z-disk disintegration and aggregate formation [42]. Muscle specific functions of STK38 were initially attributed to the regulation of the RNA-binding protein RBM24 [54]. However, based on the findings reported here, we would propose that dysregulated disposal of Z-disk components significantly contributes to the collapse of the sarcomeric architecture observed upon loss of STK38. Indeed, similar to the accelerated turnover of filamin and SYNPO2 demonstrated by us in STK38-depleted smooth muscle cells (see Fig. 2), knockdown of the kinase in cardiac muscle cells stimulated the degradation of several Z-disk and cytoskeleton proteins [42]. Apparently, STK38 fulfills a central role in the balancing of anabolic and catabolic processes in muscle cells.

## Acknowledgements

We thank Karen Himmelberg for expert technical assistance and Dieter Fürst for providing antibodies against cytoskeletal proteins. Work was supported by the following grants: DFG EXC 1010 SyNergy and support by the Boehringer Ingelheim Foundation to C.B., Wellcome Trust Research Career Development fellowship 090090/Z/09/Z to A.H., and DFG FOR 1352 TP10, DFG FOR 2743 TP1 and DFG HO 1518/9-1 to J.H.

## Authors contributions

C.K., J.W., R.J., B.K., D.S. and J.H. conducted the experiments.

C.B. supervised the proteomics part of the study.

C.B., A.H. and J.H. designed the experiments, supervised the involved coworkers and discussed the obtained data.

J.H. wrote the manuscript.

C.K., C.B. and A.H. edited the manuscript.

---

**Fig. S1.**
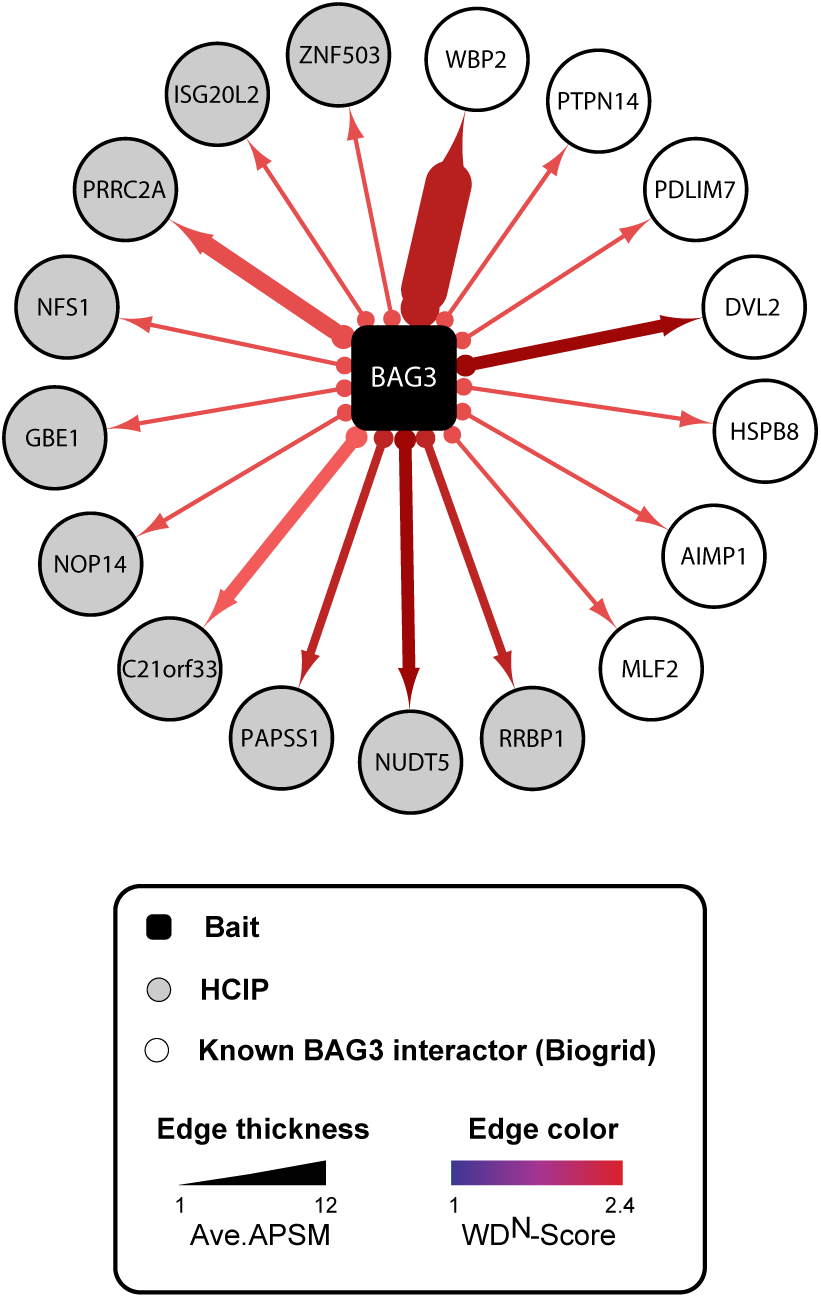
BAG3 interaction network. Lysates of 293T cells stably expressing HA-tagged BAG3 were subjected to HA-IP, followed by trypsin digestion and mass spectrometric analysis. High-confidence candidate interacting proteins (HCIPs) are color-coded according to WDN-score. Line thickness indicates the relative abundance (as average peptide spectral matches [APSMs]) of HCIPs in BAG3 immunoprecipitates. Open circles refer to known BAG3 binding proteins based on annotation in BioGrid.

**Fig. S2.**
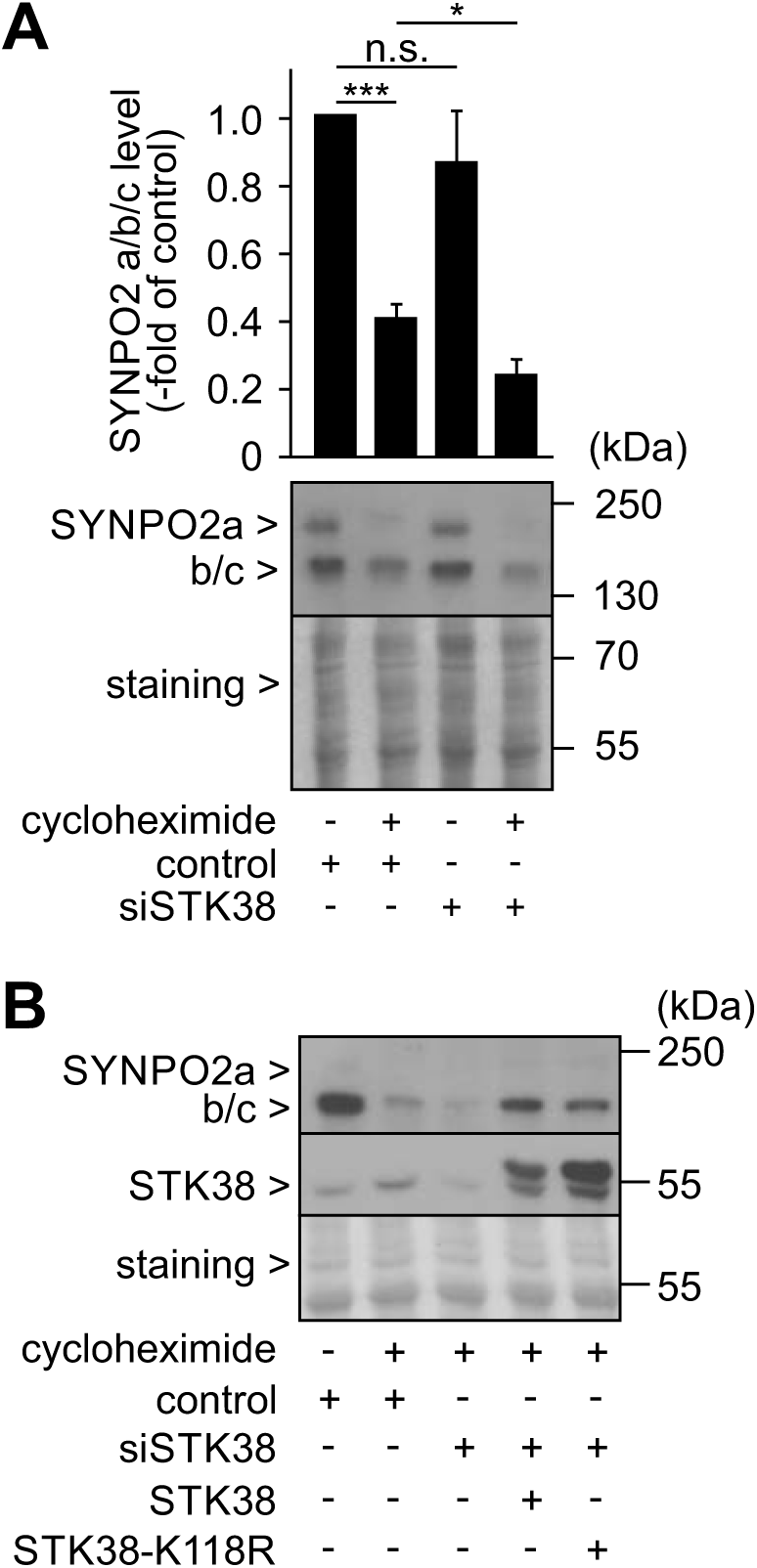
Modulation of STK38 levels in smooth muscle cells affects SYNPO2 degradation. (A) STK38 was depleted in A7r5 smooth muscle cells transiently transfected with a corresponding siRNA (siSTK38) for 48 h. Control cells received allstars negative control siRNA (control). When indicated, cells were incubated with cycloheximide (50 μM) for 3 h prior to lysis. Signal intensities were quantified and normalized to the level of total protein detected in the same sample by Ponceau S staining (staining). SYNPO2 level in control cells was set to 1. 50 μg protein were loaded per lane. Indicated proteins were detected by western blotting with corresponding antibodies. Data represent mean values +/- SEM: n = 4, *p ≤ 0.05, ***p ≤ 0.001, n.s. - non significant. (B) Rat A7r5 smooth muscle cells were transiently transfected with an siRNA against rat STK38 (siSTK38) for 48 h. Control cells received allstars negative control siRNA (control). When indicated, cells were cotransfected with plasmids encoding human STK38 or STK38-K118R, all other samples received empty plasmid. Cells were treated with cycloheximide (50 μM) for 3 h prior to lysis as indicated. 50 μg protein were loaded per lane. Indicated proteins were detected by western blotting with corresponding antibodies.

**Fig. S3.**
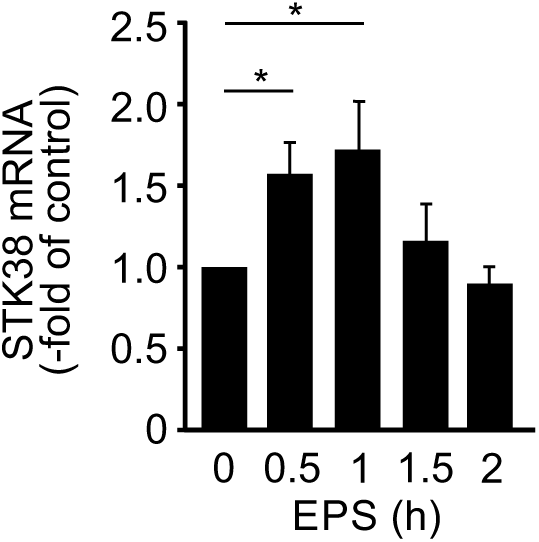
Transcription of *STK38* is stimulated in differentiated C2C12 myotubes by EPS treatment. Differentiated C2C12 myotubes were subjected to electrical pulse stimulation (EPS) for the indicated times, followed by lysis and transcript quantification by quantitative real time PCR. Transcript level in control cells was set to 1. Data represent mean values +/- SEM: n = 5, *p ≤ 0.05.

**Supplemental Table S1.**
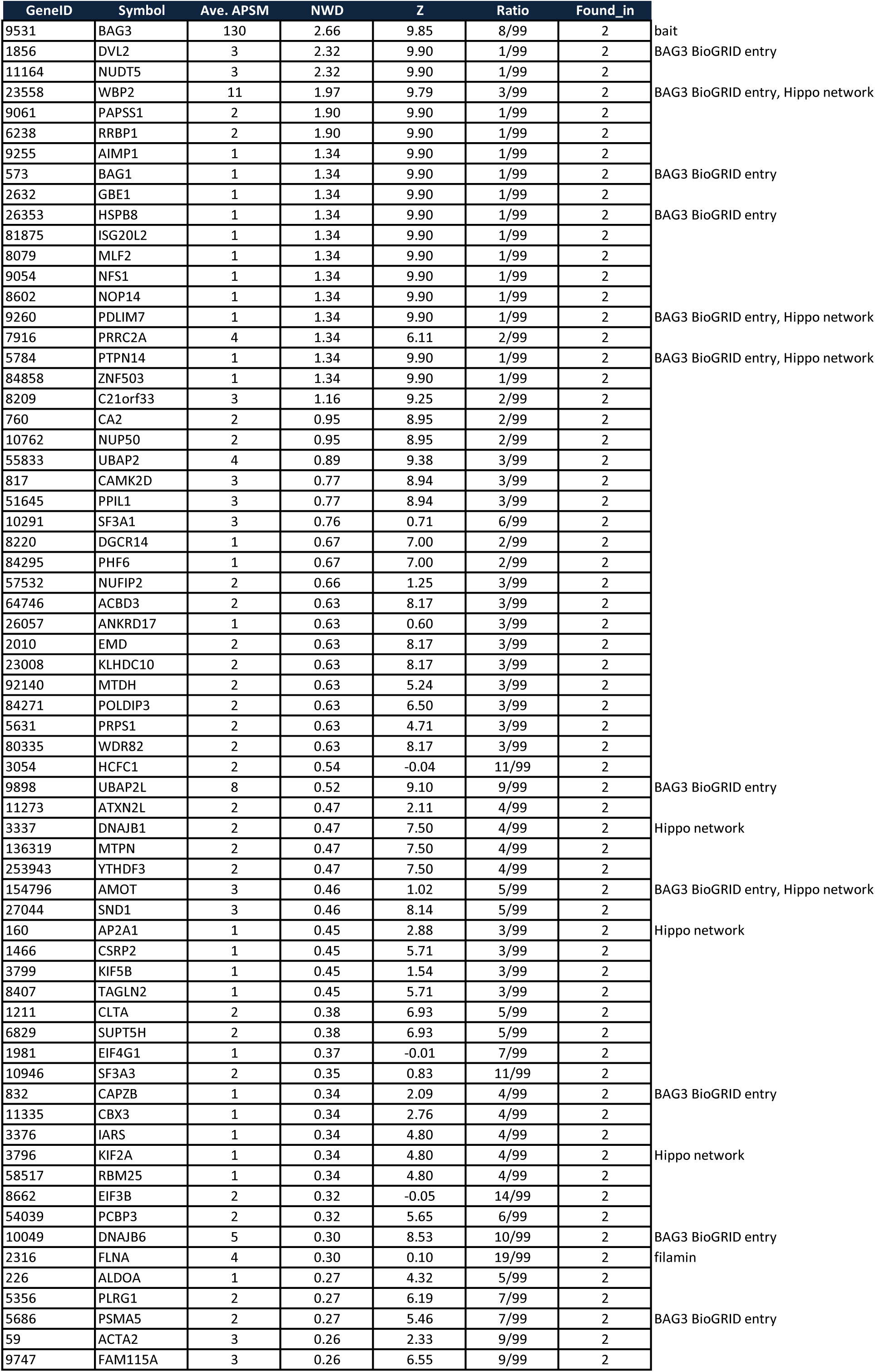
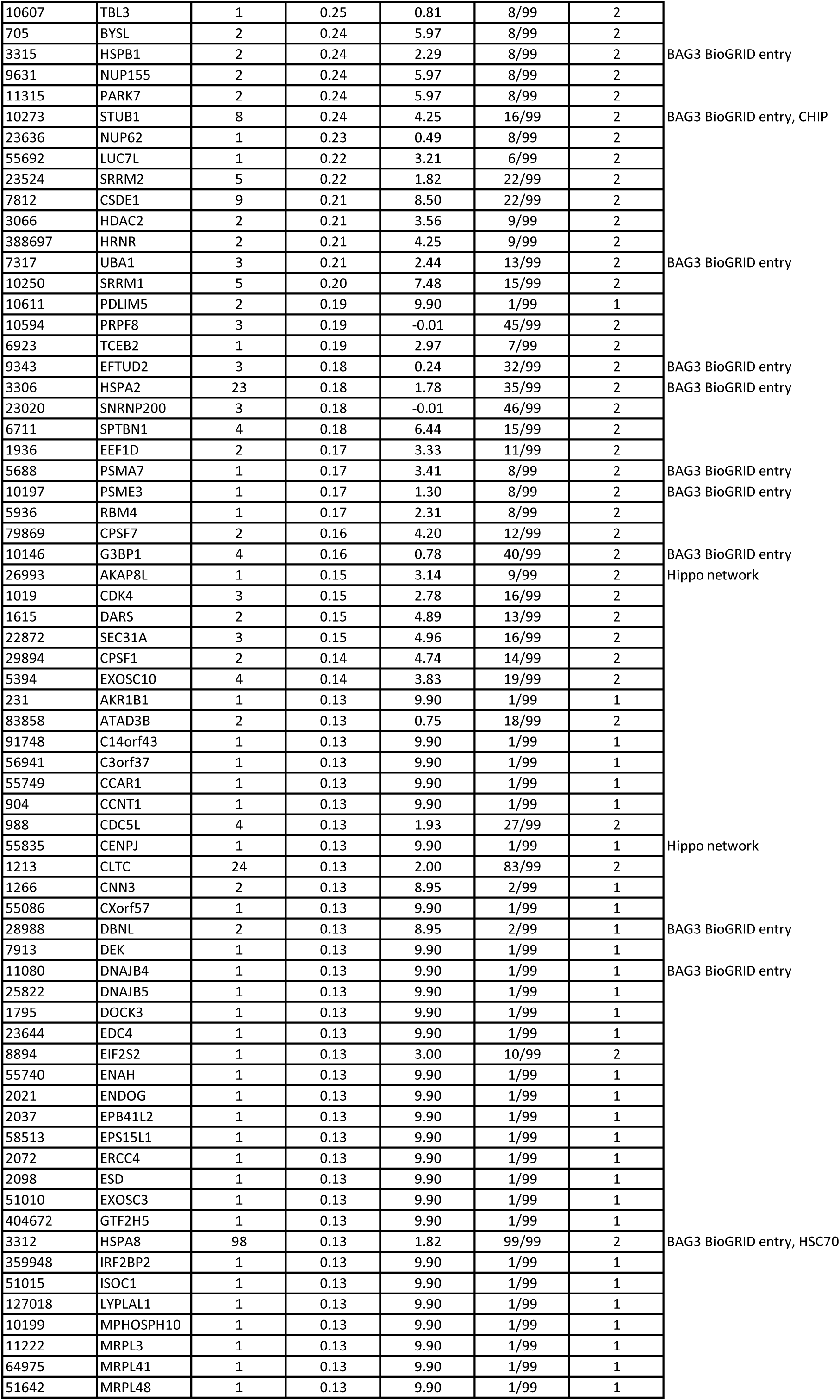
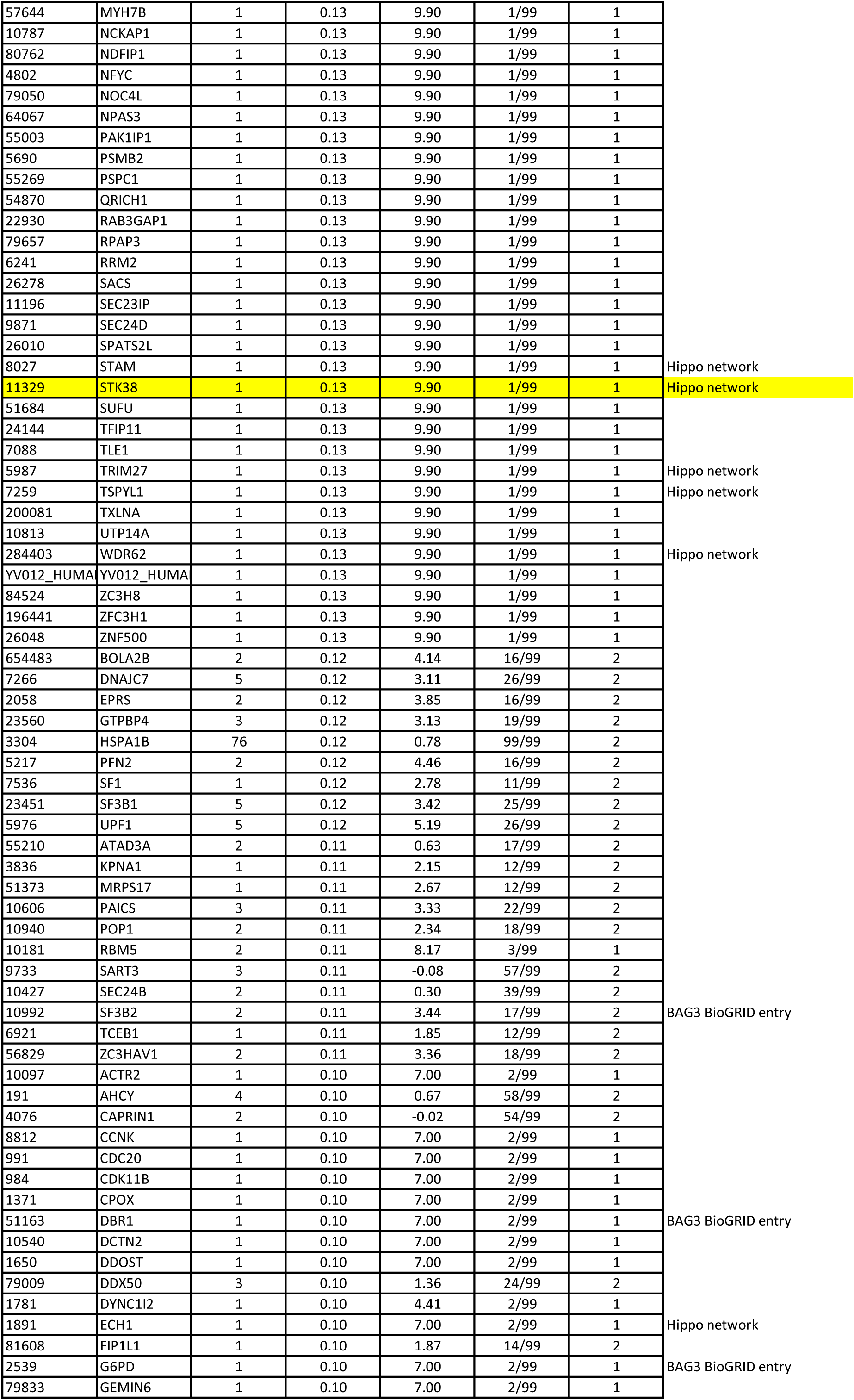
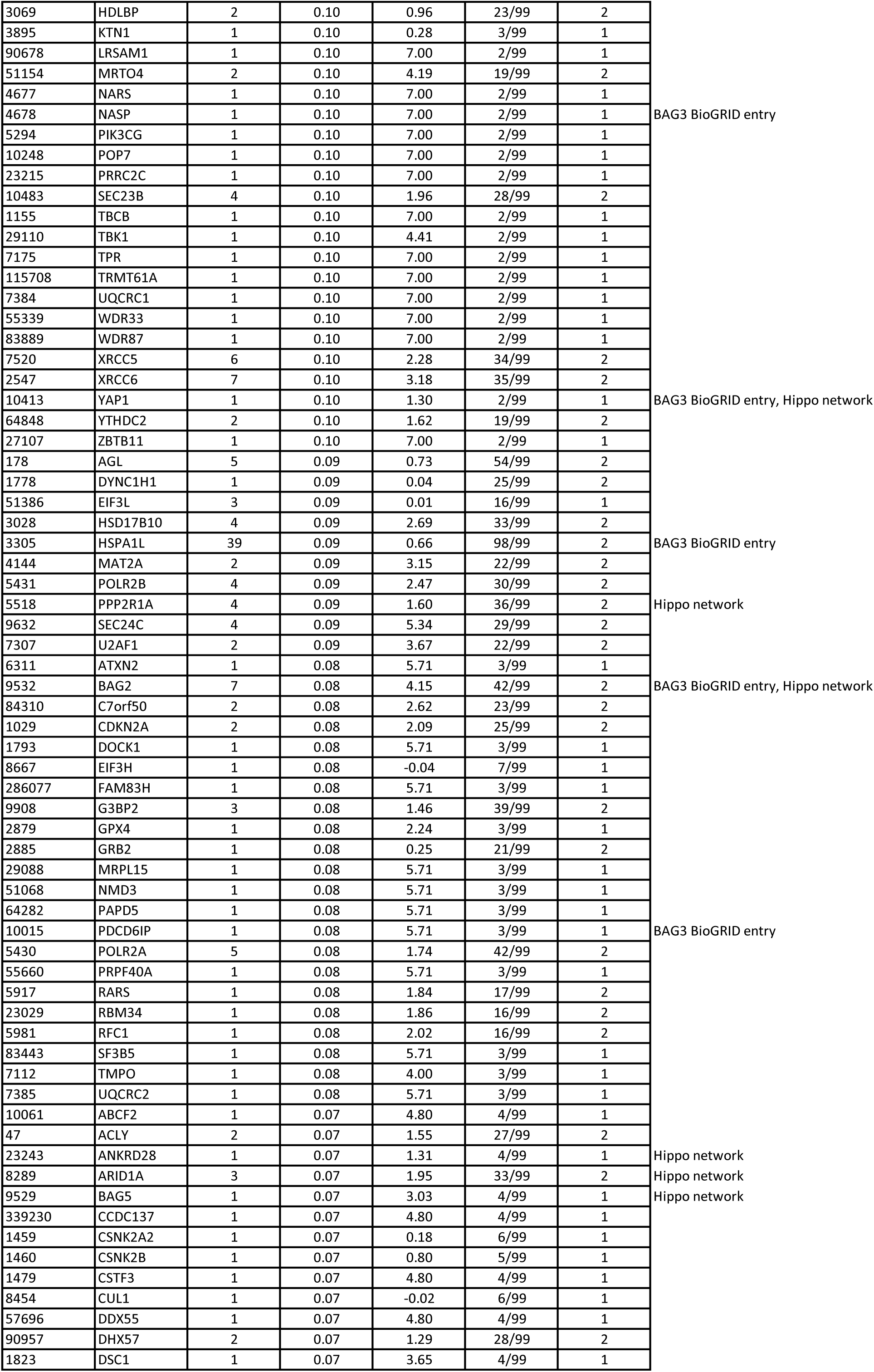
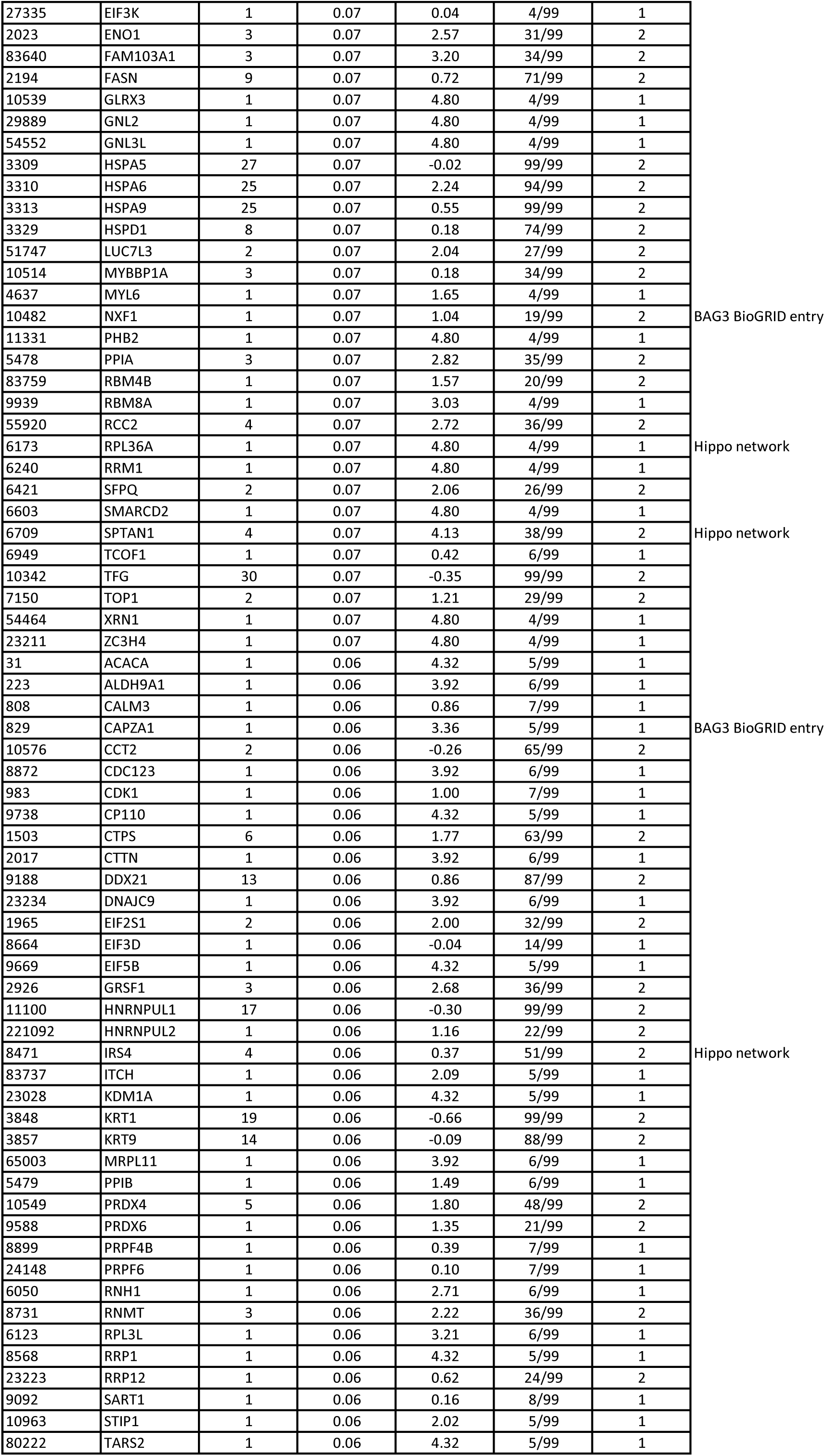
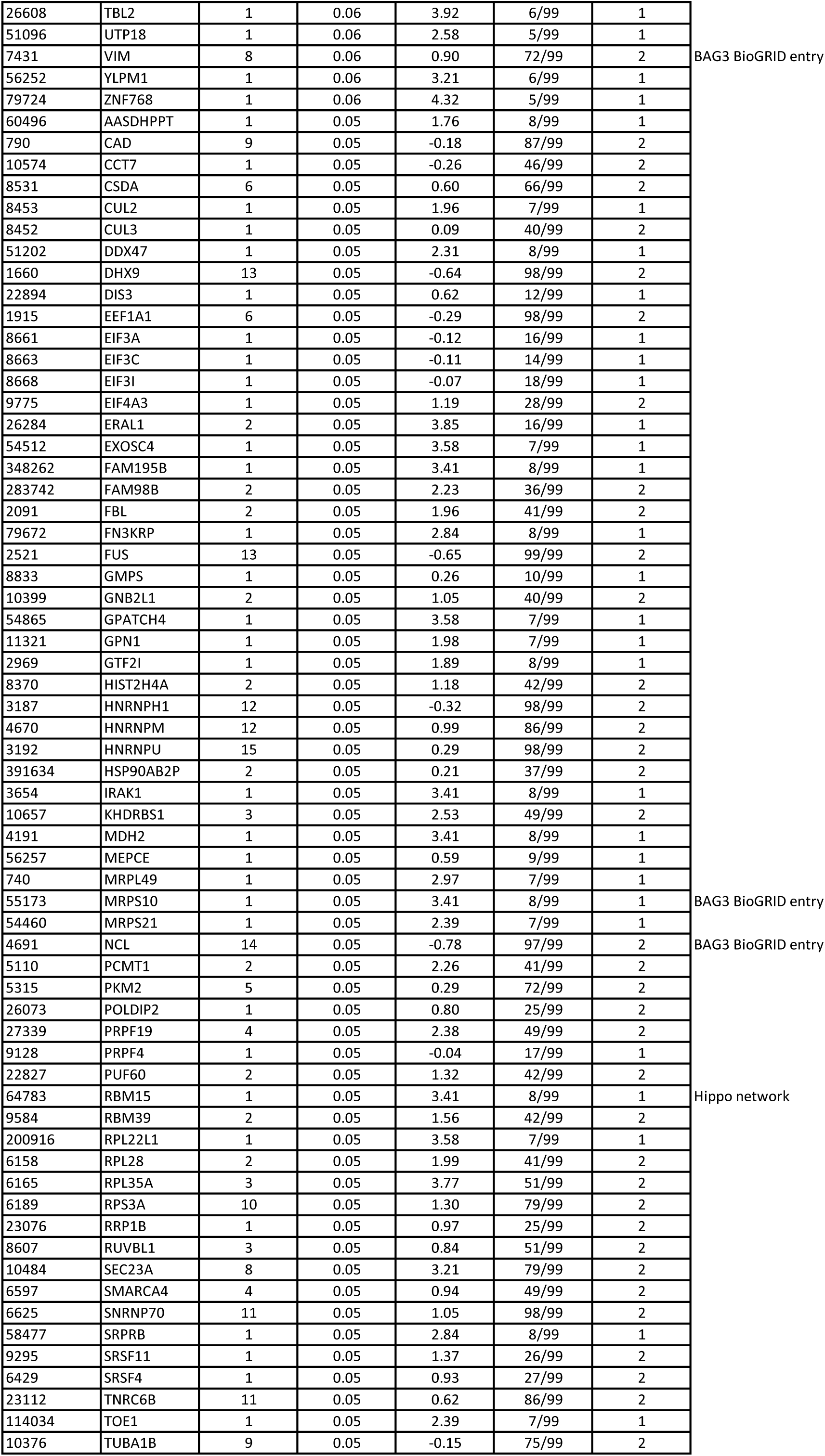
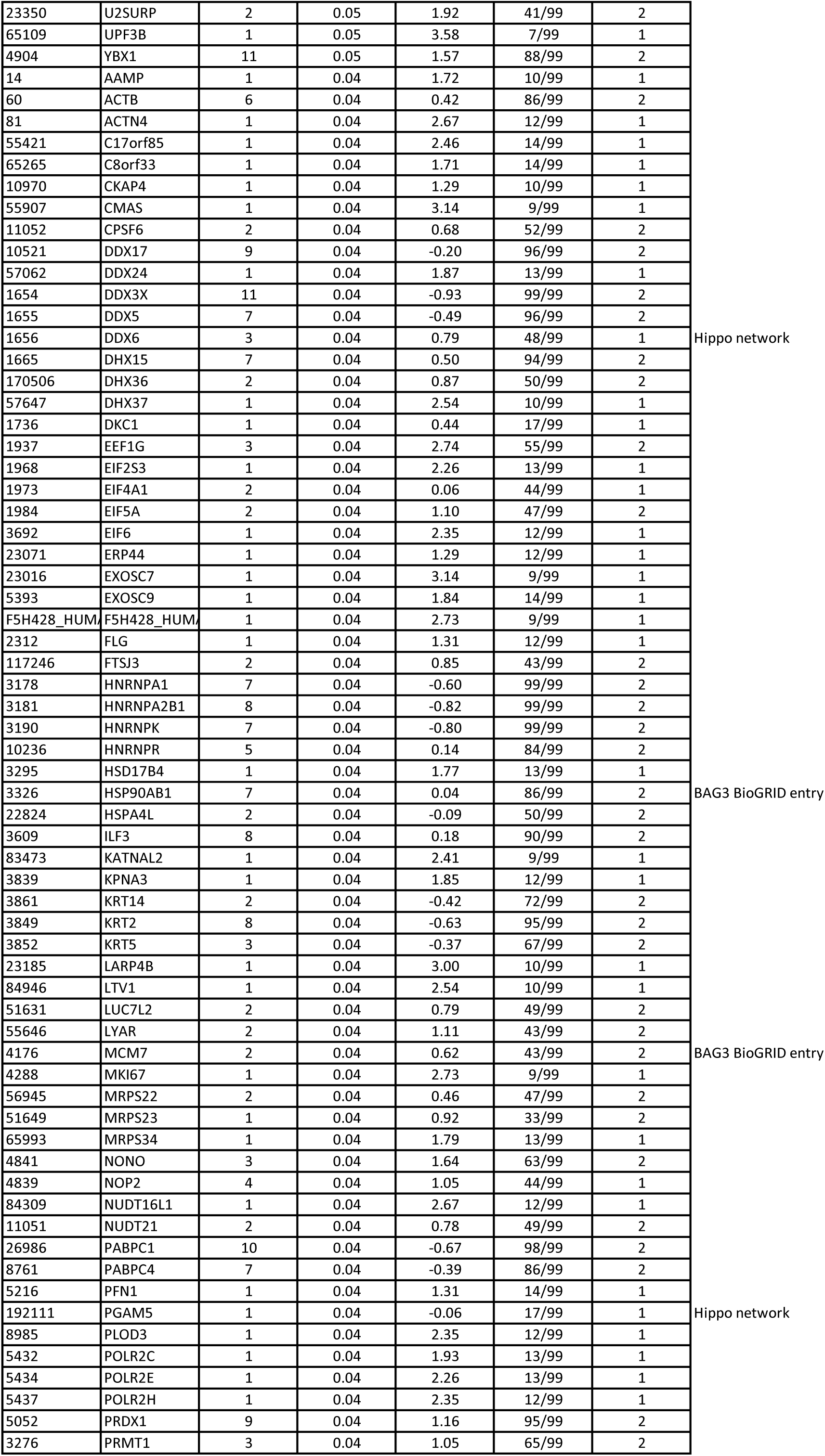
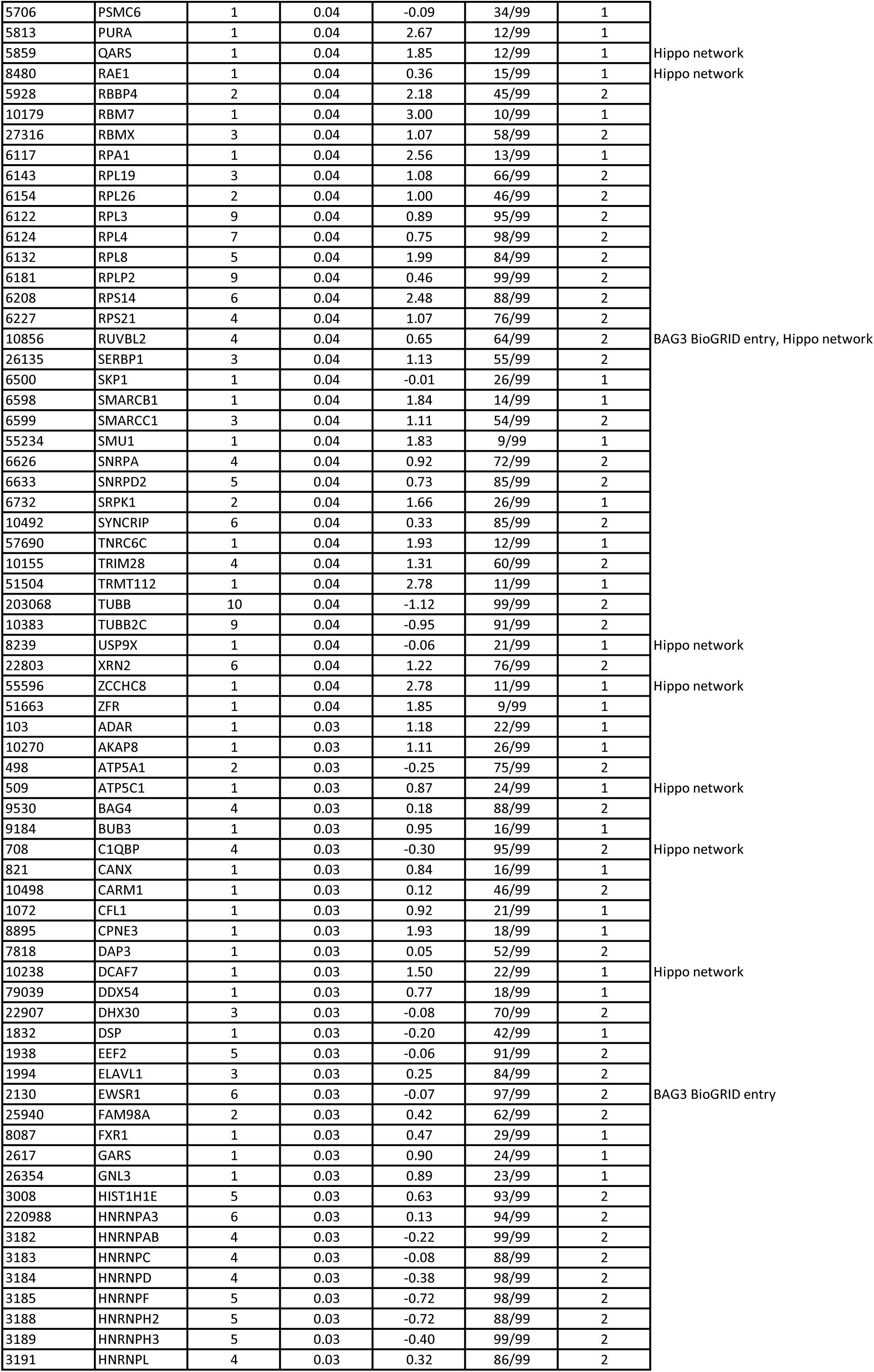
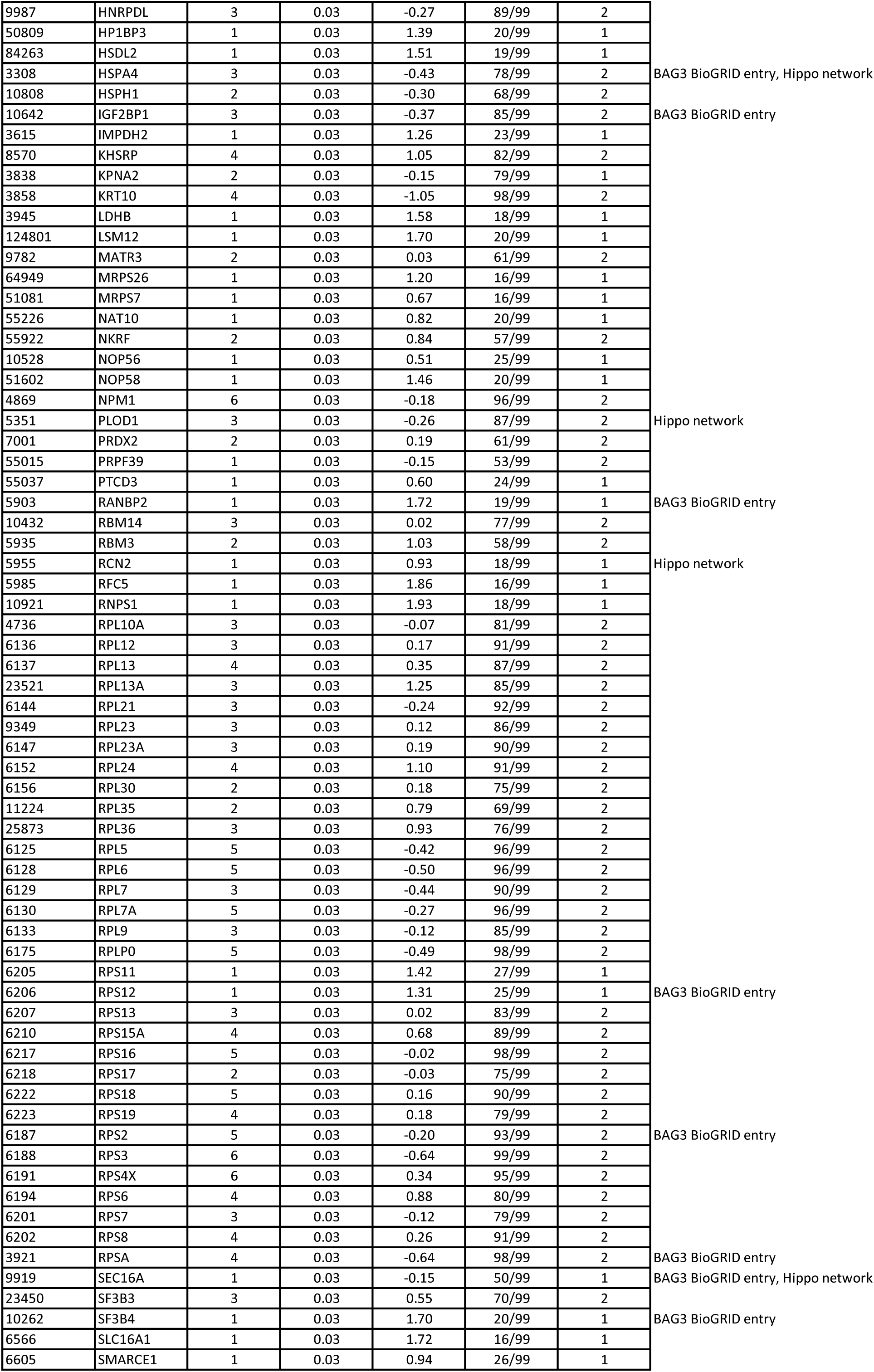
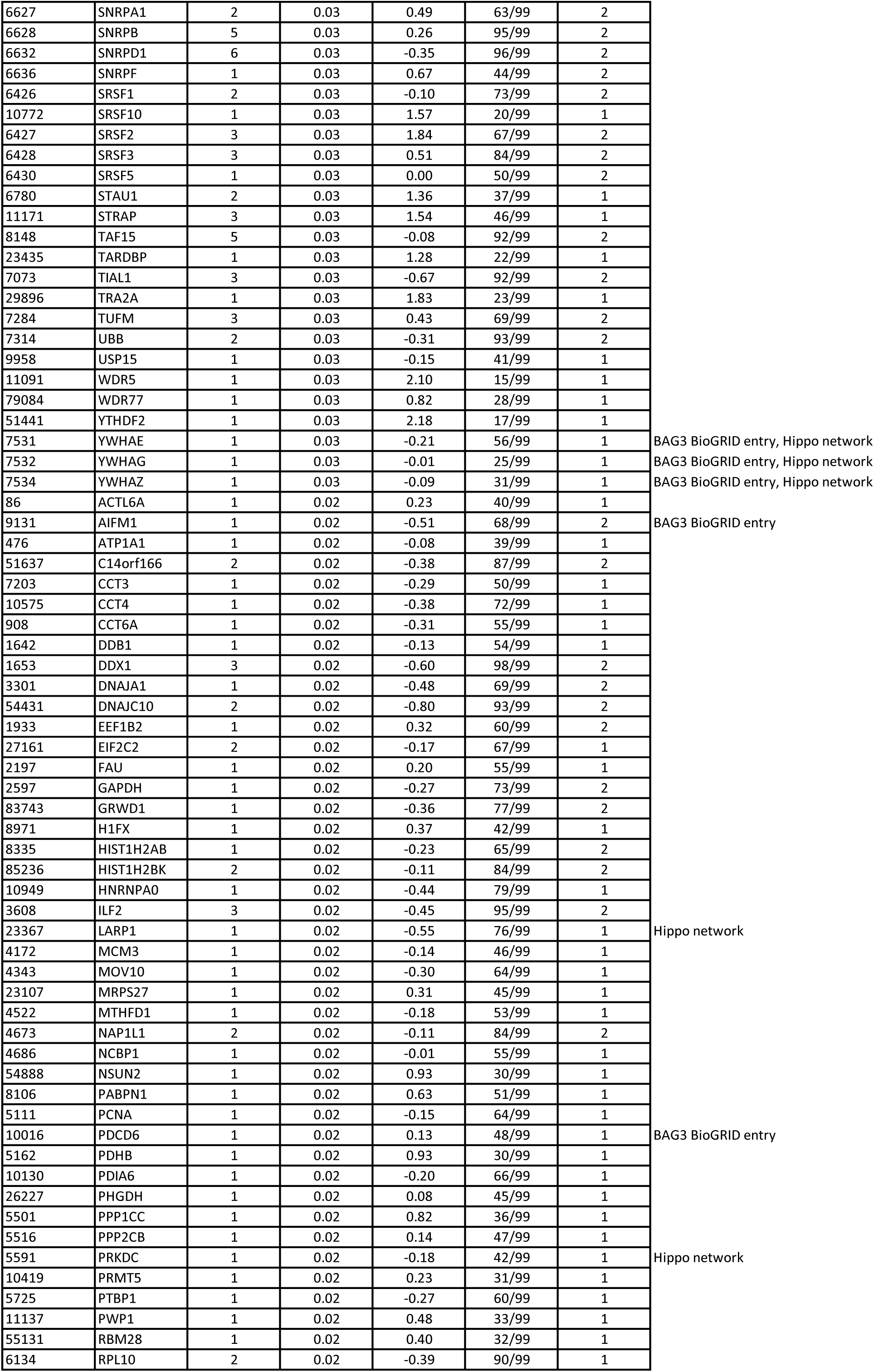
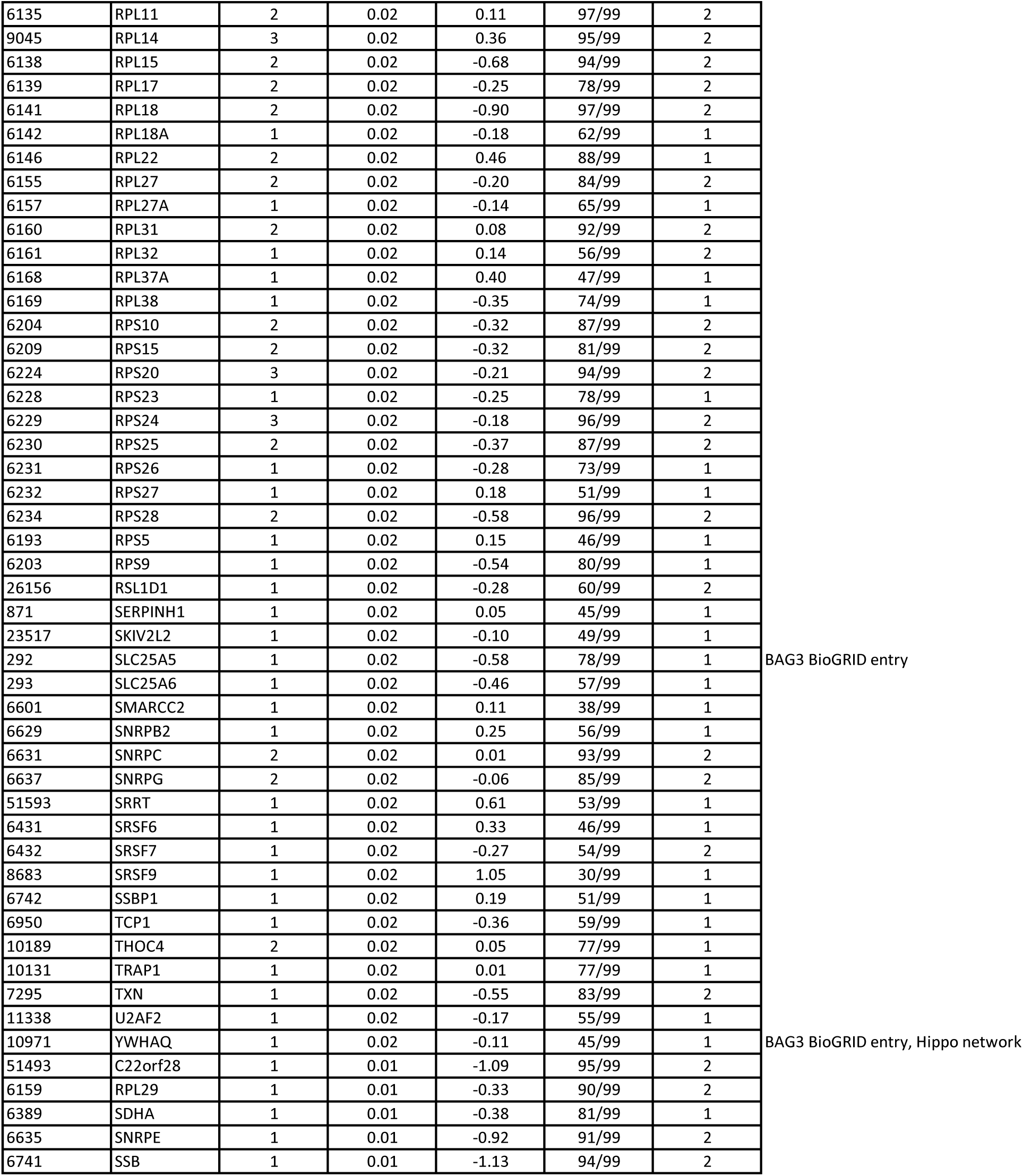
(Related to Figure 1) Proteomic characterization of BAG3 complexes isolated from HEK293T cells stably expressing N-terminally HA-tagged BAG3. APSM stands for “average protein spectral matches” and takes into account peptides which match more than one protein in the database NDW stands for „normalized weighted D (WD^N^) score” and reports the frequency, abundance and reproducibility of each interaction

